# Structure and functional analysis of CDNF-BiP interaction reveal a role in endoplasmic reticulum proteostasis regulation

**DOI:** 10.1101/2025.07.08.663636

**Authors:** Satoshi Fudo, Olesya Shpironok, Oleksii Zdorevskyi, Urve Toots, Mart Ustav, Li-Ying Yu, Mart Saarma, Tommi Kajander

## Abstract

Cerebral dopamine neurotrophic factor (CDNF) was identified for ability to rescue midbrain dopamine neurons and as endoplasmic reticulum (ER)-located protein that can be secreted. Structurally homologous to mesencephalic astrocyte-derived neurotrophic factor (MANF), CDNF has been shown to interact with ER-localized chaperone BiP. The ER plays a crucial role in protein synthesis, folding, and quality control, with BiP being a key player in maintaining protein homeostasis. CDNF is protective against ER stress and involved in regulating the unfolded protein response (UPR) signaling and interaction with BiP. Recent studies have shown that CDNF interacts with UPR sensor proteins PERK, IRE1α, and ATF6, suggesting an overlap in CDNF binding with UPR sensors and BiP. In rodent models of Parkinson’s disease (PD), CDNF protects and restores the function of brain dopamine neurons and was successful in PD phase 1 clinical studies. CDNF has shown therapeutic potential for several neurological diseases, including amyotrophic lateral sclerosis, and ischemic stroke. Despite extensive knowledge on CDNF’s impact on cellular function and neuronal degeneration, its detailed molecular mechanism of action in the ER remains unclear. Here, we have characterized the CDNF interaction with BiP both structurally and functionally and solved the crystal structures of CDNF-BiP complexes to 1.5 Å resolution, complemented with molecular dynamics simulations. Results show CDNF’s role as an antagonist of BiP nucleotide exchange, and thus in its chaperone function, binding to the ADP-bound state. Finally, we show its effect on neuroprotection with stem cell-derived human dopamine neurons, highlighting its potential in neurodegenerative disease treatment.

## 1. Introduction

Cerebral dopamine neurotrophic factor (CDNF) was originally identified for its capability to rescue midbrain dopamine neurons^1^ and later was found to be endoplasmic reticulum (ER)-located protein which can be secreted^2, 3^. It is structurally homologous to mesencephalic astrocyte-derived neurotrophic factor (MANF)^4, 5, 6^, similarly located mainly in the ER. Earlier it has been shown in cells and by proteomic analyses that CDNF interacts with ER-localized Hsp70 family chaperone BiP, also known as GRP78, and with other chaperones^7^.

The ER has a major function in protein synthesis, folding and quality control of proteins targeted outside cytoplasm. Different chaperones regulate the protein synthesis and processing in the ER and maintain the protein homeostasis. A key player in this maintenance of protein folding balance in the ER is BiP^8, 9, 10^. Several cochaperones regulate and aid the function of BiP and other Hsp70 proteins. BiP is a two-domain protein with the N-terminal nucleotide binding domain (NBD), or ATPase domain, and the C-terminal substrate binding domain (SBD)^11, 12^ (Fig. 1A). Overall, Hsp70 family proteins, including BiP, bind to unfolded proteins when they accumulate under cellular stress^13^. Binding of substrate proteins to BiP is nucleotide dependent. Several nucleotide exchange factors facilitate the turnover of ADP to ATP and further release of substrate^14^.

**Figure 1.**
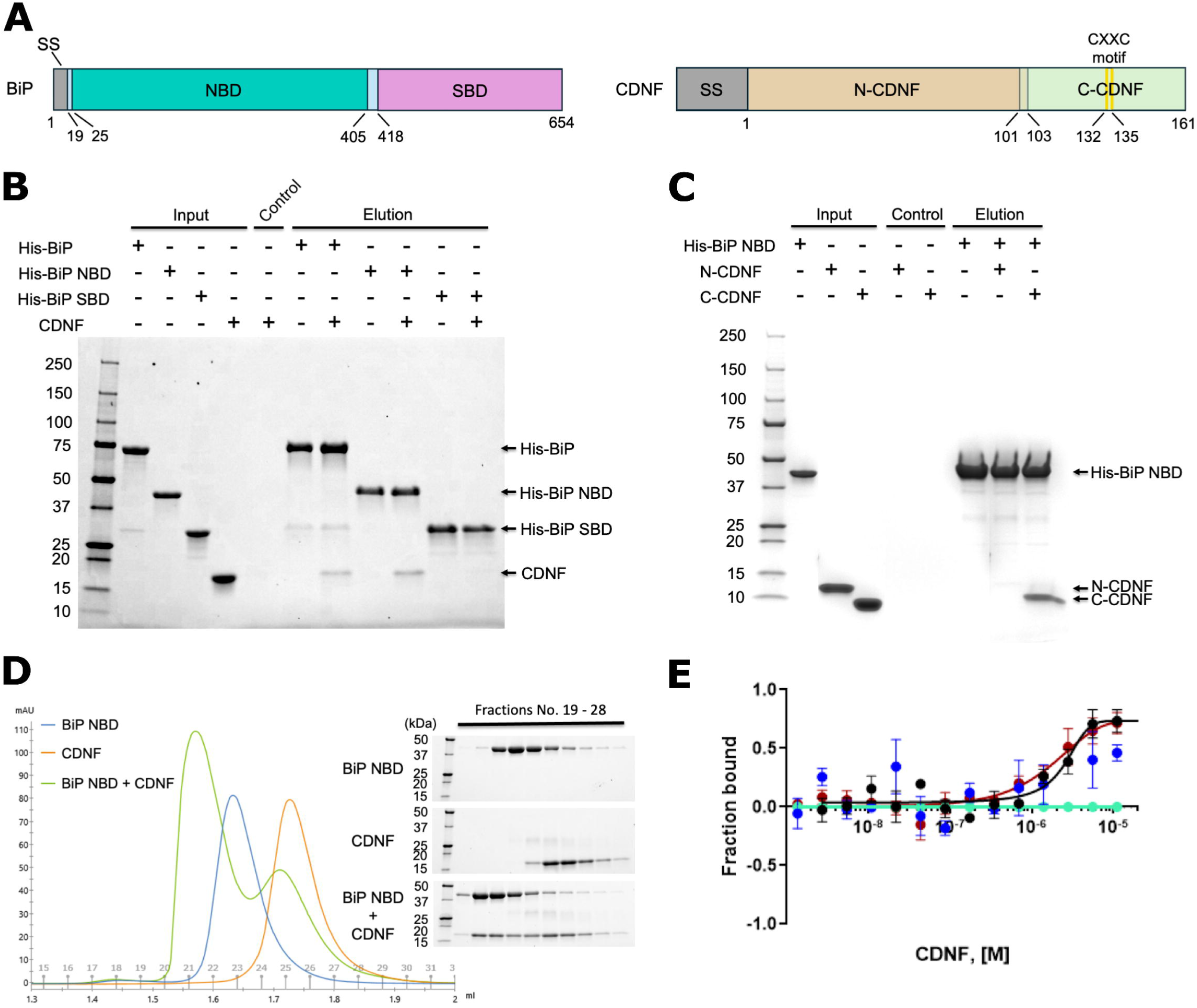
Characterization of BiP-CDNF interaction. **(A)** Schematic primary structures of human BiP and CDNF. The signal sequence (SS) of each protein is marked as grey, the nucleotide binding domain (NBD) of BiP as turquoise blue, the substrate binding domain (SBD) as pink, the N-terminal domain of CDNF (N-CDNF) as light orange, and the C-terminal domain of CDNF (C-CDNF) as pale green. The two yellow bars mark the conserved cysteine residues of the CXXC motif within C-CDNF. Before the SS is cleaved off, CDNF is 187 amino acids long, and the mature protein is 161 amino acids long. CDNF numbering starts from the N-terminus of the mature form. **(B-C)** Pull-down assays using the Ni-NTA resin. Different His-tagged BiP proteins were immobilized to the resin and different CDNF proteins were incubated. Molecular wight markers in kDa are shown on the left. **(D)** Analytical size exclusion chromatography (SEC) for BiP NBD alone (blue), CDNF alone (orange), and BiP NBD plus CDNF (green) with a Superdex 200 Increase 3.2/300 column (left). Elution fractions No. 19-28 for each sample were further analyzed by SDS-PAGE (right). Fractions No. 19-28 were loaded in numerical order from left to right in the gels. **(E)** Microscale thermophoresis (MST) analysis showing the binding curve of CDNF and full-length BiP in the absence and presence of ATP or ADP. The addition of 2 mM or 4 mM ATP (blue and light green curves, respectively) abolishes the CDNF-BiP interaction, whereas 2mM ADP maintains the binding (in red), similar to apo-BiP (black).

Both CDNF and MANF are protective against ER stress and found to be involved in the regulation of unfolded protein response (UPR) signaling and BiP function^15, 16, 17, 18, 19, 20^. Recently, it was shown that MANF binds to BiP and regulates its chaperone function^21^. Both proteins appear to have a role in BiP function and in UPR, the latter of which can be either through direct interaction with UPR receptors IRE1α, PERK, and ATF6, or potentially through BiP^7, 20, 22^. Further, in rodent Parkinson’s disease (PD) models, CDNF protects and restores the function of brain dopamine neurons^1, 23^, with the latter study demonstrating efficacy in an α-synuclein-based model, and CDNF is in clinical studies for PD^24^. In line with that, CDNF knockout mice have a loss of enteric dopamine neurons and an age-dependent deficit in the function of the midbrain dopamine system, which are reminiscent of those seen in early stages of PD^18^. *In vivo*, CDNF has shown therapeutic potential towards several neurodegenerative diseases, including PD, amyotrophic lateral sclerosis (ALS), Huntington’s disease and ischemic stroke^6, 17, 25, 26^.

Despite all the knowledge on CDNF impact on cellular function and neuronal degeneration, the detailed molecular mechanism of action of CDNF in the ER remains unclear. We have recently shown that CDNF interacts with luminal domains (LD) of the UPR receptor proteins PERK, IRE1α and ATF6, PERK being the primary receptor for CDNF^22^. This, together with BiP interaction with UPR receptor proteins IRE1α and PERK^27^, suggests an overlap between UPR and BiP functional regulation by CDNF, potentially critical for understanding its molecular mechanism in cellular stress response and towards PD treatment. A recent study by Graewert et al. (2024)^28^ suggested a small-angle X-ray scattering (SAXS)-based model for the CDNF interaction with BiP at low resolution. We set to characterize the CDNF interaction with BiP in detail both structurally and functionally to understand better its role in ER stress regulation and in neuronal survival.

CDNF is an α-helical two-domain protein similar to MANF in structure^29^ (Fig. 1A). Here, we show and analyse quantitively the direct interaction of CDNF with BiP. Importantly, we show its role as an antagonist of BiP nucleotide exchange and thus in the BiP chaperone cycle regulation and report the atomic resolution crystal structures of C-terminal domain of CDNF (C-CDNF) bound to BiP NBD at 1.5 Å and 1.65 Å resolution with and without bound ADP, both in the presence of Mg^2+^ and inorganic phosphate (PO_4_^2-^). We then characterized and confirmed the role of the binding interface residues on CDNF function in BiP interaction by site-directed mutational analyses *in vitro* with purified proteins and in cellular context. To complement these experiments, we analyzed the interaction complex and effect on BiP conformation and nucleotide release by CDNF further with molecular dynamics simulations, which support our conclusions. Finally, we show that CDNF-BiP interaction has an effect on human dopamine neuron survival, coupling this interaction with the role of CDNF in UPR and cellular ER stress response.

## 2. Results

### CDNF directly interacts with BiP

We first sought to confirm the direct interaction of CDNF with BiP and then characterize the interaction in detail. Previously, Eesmaa et al. (2022) demonstrated that the proteins interact, based on affinity pull-down from cell lysate coupled to mass spectrometry, cellular interaction using bimolecular fluorescence complementation assay (BiFC), and intracellular co-localization^7^. Here, we confirmed with purified proteins that full-length CDNF binds to full-length BiP and its NBD, but not the BiP SBD, and that BiP NBD binds to full-length CDNF and C-CDNF, but not to N-terminal domain (N-CDNF), as shown by pull-down assays with his-tagged BiP proteins (Fig. 1A-C). We then performed analytical size exclusion chromatography (SEC) to confirm the complex formation by CDNF and BiP NBD (Fig. 1D). At 100 µM protein concentration, we were able to observe shift of the chromatography peak indicating the presence of the complex and verified it by SDS-PAGE from the eluted peak fractions the. Microscale thermophoresis (MST) was then used to measure the affinity (*K*_d_) between BiP NBD and CDNF, which was shown to be 1.04 ± 0.43 μM (Fig. 1E) and confirmed by surface plasmon resonance (SPR) (Supplementary Fig. S1, with *K*_d_ = 23.4 ± 1.4 µM). The values fall in the typical range for dynamic, transient biological interactions. MST with ATP-bound BiP did not show any affinity for CDNF, while ADP-bound BiP had similar affinity as the apo form, in keeping with the observation that conformation of BiP in the ADP-bound state is similar to that of the apo state, while the ATP-bound state has a completely different conformation (Supplementary Fig S2). Therefore, we conclude that CDNF prefers the ADP-bound conformation of BiP and is unable to bind to the ATP-bound conformation (Fig. 1E, Supplementary Fig S2). In all cases Mg^2+^ was present in the solution, as this is presumed to be required for nucleotide binding.

### Structures of C-CDNF in complex with BiP NBD

To obtain high resolution information on the CDNF-BiP interaction, we crystallized C-CDNF with BiP NBD. Despite attempts, no crystals were obtained for full-length CDNF in complex with BiP. Notably, previously no crystal structure of C-CDNF part of CDNF had been solved, as earlier only the N-CDNF was crystallized^29^ (Fig. 1A). Thus, we here for the first time show the high-resolution structure of the C-CDNF (Fig. 2), in addition to revealing the interaction complex structure with BiP NBD. Crystal structures of the complex were solved to 1.5 Å and 1.65 Å with and without bound ADP, respectively (Fig. 2, Supplementary Table S1). In both complex structures, BiP NBD has bound Mg^2+^ and PO_4_^2-^ ions at the nucleotide binding site. Bound Mg^2+^ is coordinated by the PO_4_^2-^ ion and five water molecules in the structure without bound ADP, while one of the water molecules is replaced with the β-phosphate moiety of ADP in the ADP-bound structure. We could not obtain crystals with full-length CDNF with BiP, but analysis of interaction with N-CDNF by our pull-down experiments indicates that the contribution of the N-terminal domain to the binding is negligible, and binding is driven by the C-CDNF (Fig. 1, Fig. 2). It should be noted that the two domains of CDNF have considerable freedom of orientation relative to each other due to the relatively flexible linker connecting the two domains according to the previous NMR studies^30^ and our recent SAXS analyses^31^. In the case of MANF bound to BiP, similar binding mode was reported^21^, and based on mutational studies, N-terminal domain of MANF did not contribute significantly to the binding. The results overall support similar interaction mode for CDNF as in the case of MANF, although CDNF and MANF are different proteins with dramatically different knockout mice phenotypes and could have different modes of interaction. In the BiP-MANF complex structures, no nucleotides or other ligands are present, while here we present the structure of the physiologically relevant ADP-bound state of BiP, to which CDNF and MANF are assumed to be bound in the chaperone cycle. We observe that the ADP-bound BiP NBD conformation in complex with CDNF is very similar to the BiP NBD alone with or without ADP.

**Figure 2.**
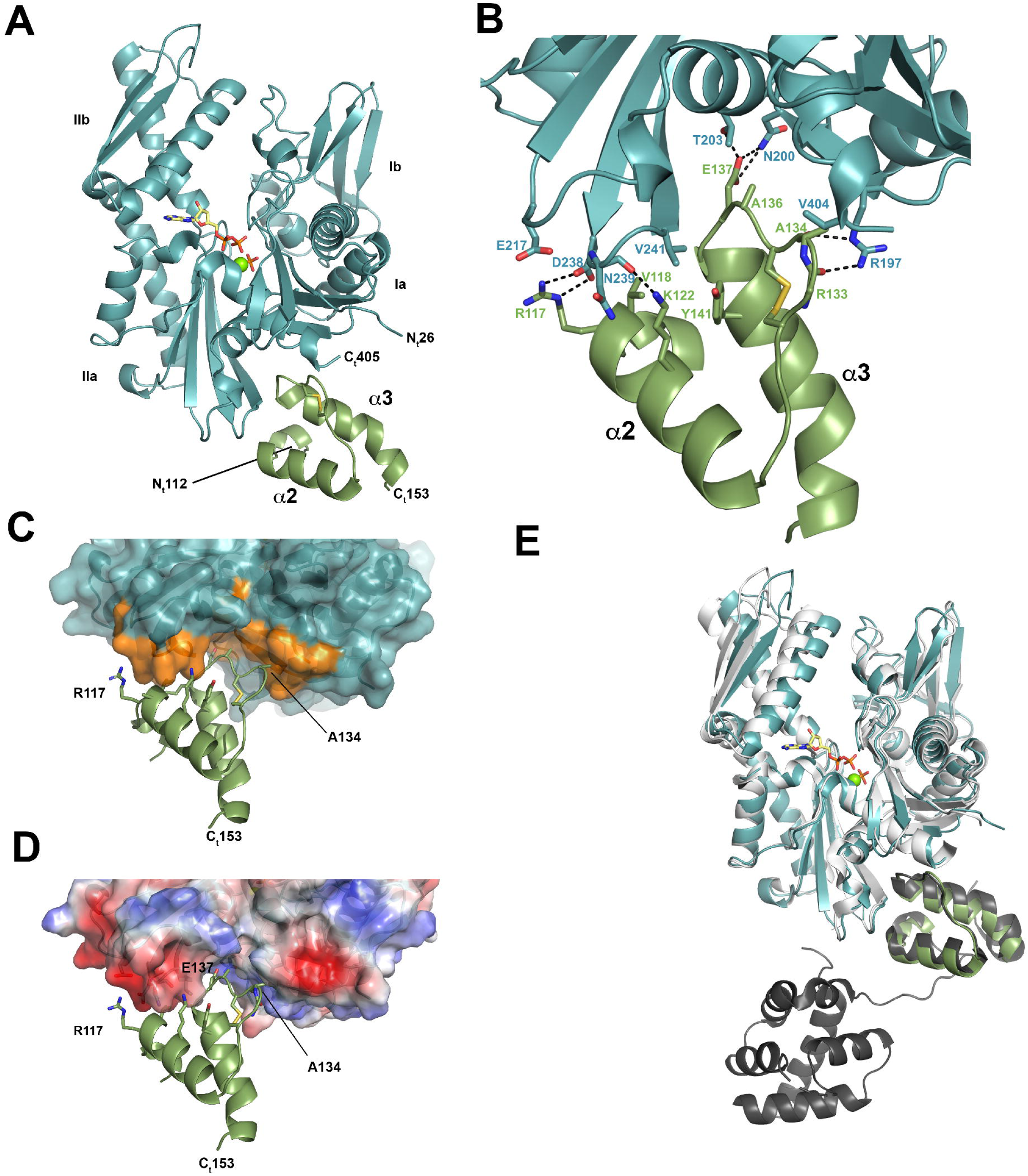
Structure of CDNF-BiP complex. **(A)** The overall structure of the C-CDNF-BiP NBD with BiP NBD in blue and C-CDNF in green. The lobes of the BiP NBD are labeled according to standard nomenclature as Ia, Ib, IIa, and IIb, and N- and C-termini are marked with “N_t_” and “C_t_” and their residue numbers. The bound ATP and P_i_ are shown as sticks, bound Mg^2+^ as a bright green sphere, and C-CDNF disulphide shown with sticks (sulphur in yellow; carbon in green), the main α-helices on C-CDNF are labeled. **(B)** The binding interface of the complex with residues and hydrogen bonds indicated. Colours are as in (A), oxygen atoms in red, nitrogen in blue, and carbon atoms follow the chain colour. Black dashed lines indicate hydrogen bonds and salt bridges. **(C)** Surface representation image of the interface with buried interface residues on BiP colored in orange (total surface area ca. 560 Å^2^). Key residues on CDNF are indicated and labeled. **(D)** Electrostatic potential displayed in BiP surface to show the charge complementarity with CDNF charged residues (including R117, K122 and E137). For full list of interface residue, please see Supplementary Table S1. **(E)** Comparison to BiP NBD-MANF structure without bound ligands, MANF (black) and BiP (grey) aligned to our structure.

The interface between CDNF and BiP has several hydrogen bonded interactions with contact surface area of 558 and 564 Å^2^ for the structures without and with ADP, respectively (Fig. 2B-D, Supplementary Table S2). There are only minor differences between the overall conformations of the ADP-bound and unbound BiP C-CDNF complex structures with overall r.m.s.d. of 0.21 Å. No significant difference was seen in the BiP-CDNF binding interface. Therefore, only ADP-bound BiP-CDNF interface is discussed further here.

CDNF docks between the Ia and IIa subdomains of BiP, interacting with BiP α-helix 200-210 between these lobes and with the tip of the IIa subdomain and through hydrophobic interface to subdomain Ia (Fig. 2A). There are two residues in CDNF that seem to play an important role in BiP binding through ionic hydrogen bonded interactions: R117 and E137 (Fig. 2B-D, Supplementary Tables S2, S3). CDNF R117 makes a bidentate hydrogen bonded salt bridge with BiP D238, while CDNF E137 makes hydrogen bonds with BiP N200 and T203. Other CDNF residues at the interface area include V118, K122, A134, A136, and Y141 (Fig. 2B). CDNF V118 is almost completely buried in the interface and forms hydrophobic interaction to BiP V241 together with CDNF Y141. CDNF has a CXXC motif including C132 and C135 forming a disulphide bridge in the C-CDNF. The loop G129-E137 between CDNF α-helices 118-128 and 138-152 stabilized by this motif is also at the binding interface, and A134 and A136 in the G129-E137 loop pack against a pocket on BiP including V404 and the C-terminal α-helix of the BiP. CDNF K122 interacts with the main chain oxygen of BiP N239 but considering only one hydrogen bonded interaction to N239 carbonyl group, this interaction is unlikely to be very strong. Indeed, K122A mutant was able to bind BiP as shown by Kovaleva et al. (2024)^22^. Similarly, this interaction was not critical for MANF interaction^21^. Comparison of the BiP-CDNF binding mode with the reported BiP-MANF shows that they have many interacting residues in common, except that the corresponding residue to CDNF A134 is G150 in MANF (Fig. 2E), and thus there is an additional hydrophobic residue in CDNF contributing to the interaction with BiP. The interface for both complexes has an ionic/hydrogen bonded component as well as a hydrophobic component, as observed from the CDNF structure. In the MANF structure by Yan et al. (2019), the ionic interactions were considered and analyzed^21^. Here, for the CDNF-BiP complex, we observe the importance of the C132-C135 disulphide bridge-containing CXXC-motif loop G129-E137 in forming a hydrophobic interface and the bidentate hydrogen bonded interaction with R197 of BiP through the CDNF backbone carbonyl groups (Fig. 2B).

### Comparison with the structures of the ADP and ATP bound states of BiP

Binding of C-CDNF seems to have little influence on the conformation of the nucleotide binding site, and the overall BiP NBD structure when compared with a known ADP-bound BiP NBD structure (PDB ID: 5EVZ)^32^ is highly similar (r.m.s.d. over Cα-atoms 0,567 Å, Supplementary Fig. S2), while the ATP binding to BiP NBD induces a large conformational change in BiP SBD^11^, and ATP-bound state is not accessible when CDNF is bound to BiP NBD, as it overlaps with the BiP linker region in the ATP-bound state of the full length protein. Thus, CDNF can only bind to BiP after ATP is hydrolyzed, as the linker region between NBD and SBD in the ATP-bound state of BiP is located in the same cleft where the CDNF binding site is located (Supplementary Fig. S2), as directly also shown by our binding assays (Fig. 1E).

### Mutational analysis of CDNF-BiP interface residues effect on binding affinity

The identified key interface residues (R117, E137, A134 and V118) on CDNF were mutated (Fig. 3A), and mutants R117E, E137A, A134Q, and V118Y and a double mutant E137A- V118Y were tested in the pull-down binding assay with BiP, where none of the mutants could be detected as binding and co-eluting with BiP (Fig. 3B). Binding of BiP NBD to the mutant CDNF proteins was also measured by MST using BiP NBD fluorescently labelled via an NHS ester-based labelling and titrated with unlabelled CDNF or CDNF mutants. All tested mutants except R117E showed no measurable affinity in the MST experiments (Fig. 3C). K122A was shown already before to bind BiP similar to WT CDNF (Kovaleva et al. 2024)^22^, which is in line with Yan et al. (2019) observations on MANF-BiP binding^21^. To complement the MST measurements, we further confirmed the inability of the CDNF point mutants to bind to BiP NBD by SPR, where all mutants showed significantly reduced binding level compared to wild type (WT) CDNF (Fig. 3D, E).

**Figure 3.**
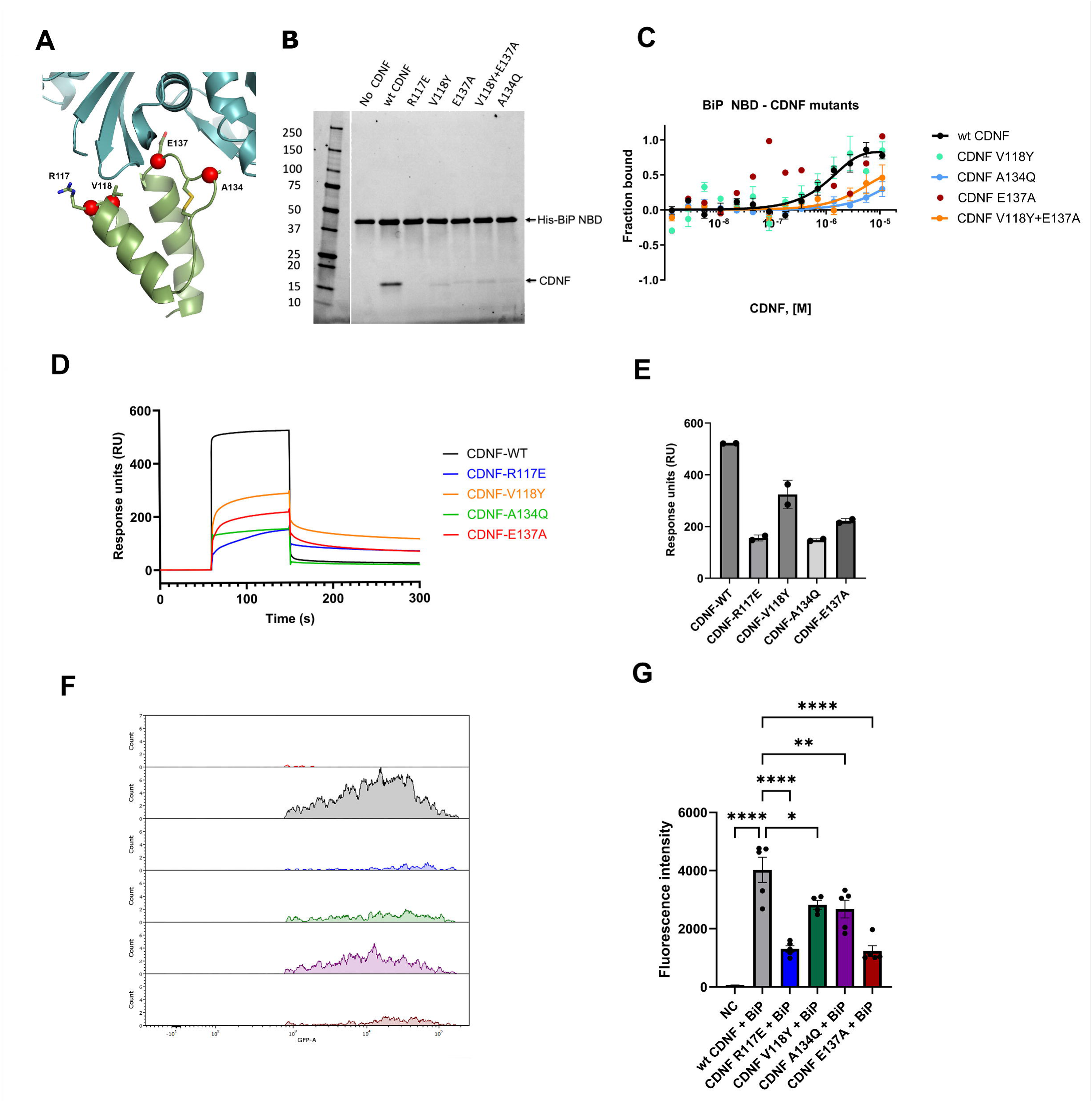
Mutational characterization of the CDNF-BiP interaction. **(A)** CDNF mutated residues shown on the complex structure highlighted with red spheres. **(B)** Pull down assay results of CDNF and the CDNF E137A, A134Q, V118Y, R117E, or V118Y-E137A mutants binding to His-tagged BiP NBD analyzed by SDS-PAGE. Molecular weight markers in kDa are shown on the left. **(C)** MST analysis of purified recombinant BiP NBD (0–5.1 µM) interaction with CDNF and with the E137A (red), A134Q (light blue), V118Y (green), or V118Y-E137A (orange) mutants. Affinity for the interface mutants was too low to be determined in the assay (*K*_d_ > 10 μM) (A134Q and V118Y-E137A) or signal could not be detected above noise (E137A and V118Y). Data shown are representative of n = 3 independent experiments. **(D)** Surface plasmon resonance (SPR) runs for analytes at a high concentration (50 μM CDNF and its mutants) binding to surface-coupled BiP NBD (see also Supplementary Fig. S1 for WT binding concentration series). Data shown are representatives of n = 2 independent experiments. **(E)** Steady-state binding levels for CDNF and its mutants analyzed from SPR in (D), indicating clearly reduced binding by the interface mutants (R117E, V118Y, A134Q and E137A). Data are presented as mean ± SD, n = 2. **(F)**. The panel shows histograms of GFP fluorescence (GFP-A) from a gated cell population used for BiFC quantification, measured by flow cytometry. The reconstituted GFP signal indicates proximity and interaction between CDNF and BiP. The negative control (red) establishes the baseline fluorescence. Cells co-transfected with WT CDNF and BiP (gray) analyzed by flow cytometry exhibit a strong GFP+ signal, indicating robust interaction. In contrast, CDNF mutants R117E (blue), V118Y (green), A134Q (purple), and E137A (brown) display reduced GFP signal intensity to varying degrees, indicating altered or disrupted interaction with BiP. These gated histograms highlight differences in BiFC signal that correspond to the mean fluorescence intensity (MFI) values used for quantitative analysis in downstream panels. Representative results from n = 3–5 independent experiments are shown. **(G)** Quantification of fluorescence intensity in BiFC assays with split-Venus fusion constructs of CDNF and BiP. Cells expressing wild-type CDNF and CDNF mutants were co-transfected with BiP and analyzed by flow cytometry. Fluorescence intensities were measured in 10,000 live cells using an LSRFortessa flow cytometer (100 µm nozzle, 488 nm excitation laser, GFP channel, 527/32 nm bandpass filter). Mean ± SD; n = 3-5, ordinary one-way ANOVA with Dunnett’s test. *p < 0.05, **p < 0.01, ***p < 0.001, ****p < 0.0001.

Importantly, also the BiFC measurements on CDNF-BiP binding with split-Venus fluorescent protein (Supplementary Fig. S3) in cellular context clearly demonstrate reduced amount of interaction for all the measured CDNF mutants compared to WT as analyzed by flow cytometry (Fig. 3F, G). Hence, in summary, the mutated CDNF residues at the binding interface clearly are important for the specific interaction with BiP, not only by electrostatic interactions but also through disruption of the hydrophobic binding pockets and the loop involving the CXXC motif including C132 and C135 in CDNF, as indicated above. The expression of CDNF mutants at a level comparable to WT CDNF and their similar purification behaviour and binding to UPR sensors strongly suggest that all tested CDNF mutants are properly folded.

### CDNF antagonizes the nucleotide exchange by BiP

We then tested whether CDNF has an effect on the nucleotide exchange by BiP. We tested the effect on nucleotide exchange with the fluorescent ADP release assay using *N*-8-(4-*N′*-methylanthranylaminobutyl)-8-aminoadenosine diphosphate (MABA-ADP) as the substrate. With the BiP NBD protein, wild-type CDNF and C-CDNF inhibited the MABA-ADP release, whereas N-CDNF failed to slow down the process (Fig. 4A, B). All the four CDNF mutants at the BiP binding interface (R117, E137, A134, and V118) generated and tested for the effect on binding hindered the inhibitory effect compared to the WT protein. The same trend was observed with full-length BiP (Fig. 4C). The inhibitory effect of CDNF was also confirmed in the presence of Grp170, a known nucleotide exchange factor (NEF) of BiP, and with physiological concentration of Ca^2+^ (Fig. 4D). The results show that CDNF affects the nucleotide exchange function of Grp170. These results confirm that CDNF works as a nucleotide exchange inhibitor, as suggested also for MANF^21^, and the residues in the C-terminal domain, which we identified from the complex crystal structures, are important for the function, whereas the N-terminal domain does not seem to be required for the BiP regulation as suggested also by the binding studies.

**Figure 4.**
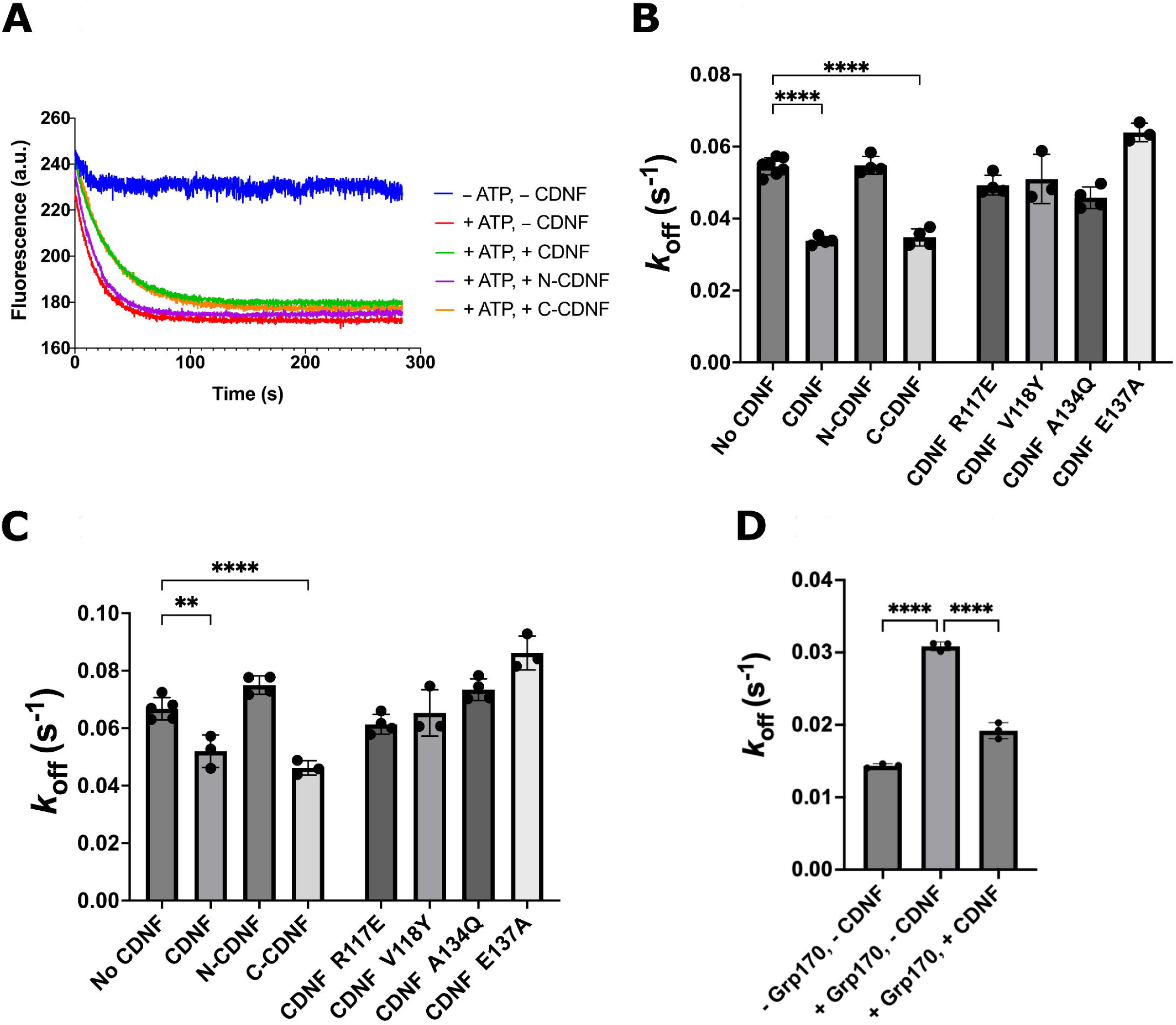
MABA-ADP release assay showing the effect of CDNF on the BiP’s nucleotide release rate. **(A)** Representative plot of fluorescence against time for each condition from BiP NBD. BiP NBD was preincubated with MABA-ADP, and solution containing ATP and CDNF (or N-CDNF or C-CDNF) was added at t = 0. **(B-D)** Mean MABA-ADP release rate constants with different CDNF constructs calculated from at least three independent experiments for BiP NBD **(B)**, full-length BiP **(C)**, and BiP NBD in the presence of physiological concentration of Ca^2+^ **(D)**. Grp170, a known nucleotide exchange factor (NEF) of BiP, was also added in (D) and the effect on nucleotide release rate was observed in the absence and presence of CDNF. For (B-D); Data are presented as mean ± SD, n = 3-7. Statistical significance was determined using ordinary one-way ANOVA with Dunnett’s test. **p < 0.01, ****p < 0.0001.

### Molecular dynamics simulations support the stabilization of ADP-bound state by CDNF

We conducted microsecond-long molecular dynamics (MD) simulations to investigate the mechanistic details of BiP nucleotide exchange inhibition by CDNF using our high-resolution structure of BiP NBD in complex with the C-terminal domain of CDNF. We observed that C-CDNF maintained a stable association with BiP NBD, where the structural hydrogen bonds (^CDNF^Glu137-^NBD^T203, ^NBD^D238-^CDNF^R117, ^NBD^R197-^CDNF^R133, see Fig. 2B) remain well-preserved throughout the trajectory (Fig. 5A). It is noteworthy that ^NBD^N239-^CDNF^K122 interaction, which was previously shown not to be critical for binding between CDNF and BiP, or MANF and BiP^21, 22^, exhibited the weakest stability among analyzed hydrogen bonds (Fig. 5A, red curve). Simultaneously, molecular contact analysis between the two subunits reveals that numerous hydrophobic interactions further stabilize the binding of C-CDNF to BiP NBD (Supplementary Fig. 4, Supplementary Table 2). The root-mean-square fluctuation (RMSF) of the backbone C-alpha carbons (see methods) revealed comparable stability of the BiP NBD in both “free” and C-CDNF-bound configurations (Supplementary Fig. S5A), with slightly higher overall RMSF values in its “free” state (Supplementary Fig. S5B). This finding is further supported by the principal component analysis (PCA) showing the overall mobility of the protein is indeed connected with the "scissors-like" reciprocal motions of the BiP NBD lobes (Fig. 5B, see also Movies S1, S2). Remarkably, structural fluctuations of lobe IIb (Fig 2A, Supplementary Fig. S6) are suppressed upon C-CDNF binding (Fig. 5B), suggesting greater flexibility of subdomain movements of BiP NBD in the absence of C-CDNF.

**Figure 5.**
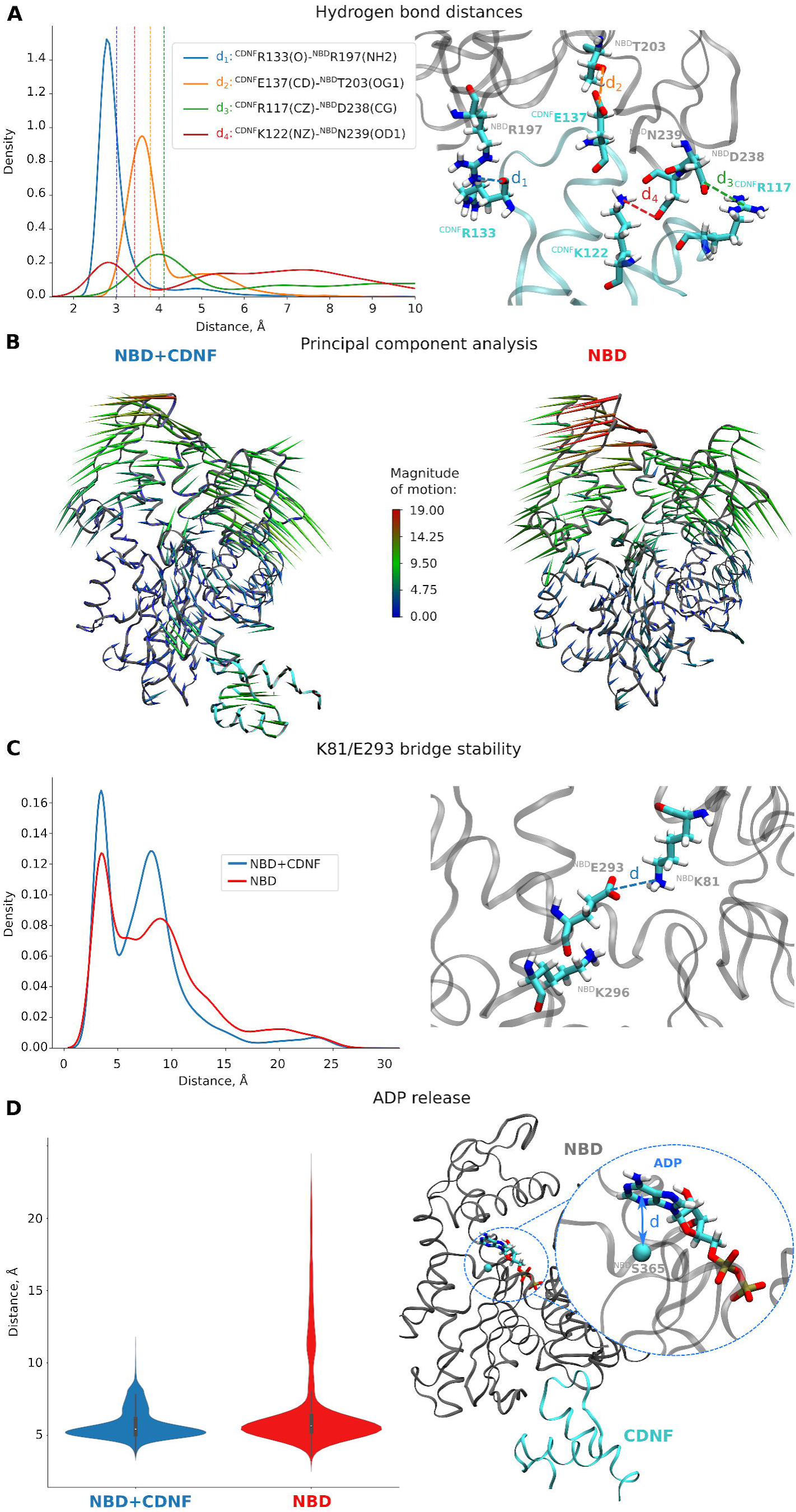
Classical MD simulations of the interactions of CDNF C-terminal domain with BiP nucleotide-binding domain. (**A) - (C)** Simulation of the high-resolution apostructure (PDBID: 9H0C). (**A)** Interactions between BiP NBD and C-CDNF subunits: analysis of hydrogen bonding residues. Vertical dashed lines on the left panel designate the corresponding distance in the crystal structure. The analyzed distances are also illustrated as dashed lines in the right inset. **(B)** Porcupine plots demonstrating protein backbone mobility along the first principal component: the slowest magnitude of motion is shown in blue, while the largest in red (see inset). The arrow lengths and colors depict the differences in the magnitude of motion: from the lowest (short) to the largest (long). **(C)** Stability of the salt bridge ^NBD^Lys81/^NBD^Glu293 sampled from our MD simulation trajectory: distribution of distances between the NZ atom of Lys and CD atom of Glu. The second peak (∼8 Å) corresponds to the competitive interaction of ^NBD^Glu293 to ^NBD^Lys296 (see inset). Blue line corresponds to the simulation of the BiP NBD + C-CDNF complex, while the red line is collected from simulation trajectories of BiP NBD alone. Smoothed distributions in (**A**, **C**) are obtained employing kernel density estimation. **(D)** Violin plots for the distances between the ADP purine ring (atoms N1- C2-N3-C4-C5-C6) and the C_α_ carbon of ^NBD^Ser365, sampled in our classical MD simulations of the nucleotide-bound structure of the C-CDNF-BiP NBD complex (see methods). The analyzed distance is illustrated in the right panel. The respective time series are depicted in Supplementary Fig. S7.

To investigate this phenomenon on a single-residue scale, we tracked the dynamics of ^NBD^K81- ^NBD^E293 salt bridge in both “free” and “C-CDNF-bound” states. In coherence with the PCA and RMSF data, the salt bridge is less present in the free state (peak ∼4 Å, Fig. 5C), thus potentiating the release of the ADP nucleotide. Next, we simulated the dynamic behavior of the ADP substrate within the nucleotide binding pocket using the high-resolution ADP-bound structure of BiP NBD in complex with C-CDNF, as determined here, and for BiP NBD alone (PDBID: 5EVZ). The results revealed an increased mobility of the ADP molecule in the BiP NBD-only setup compared to the C-CDNF-bound state. Notably, only in the absence of C-CDNF, we could observe the nucleotide release from the active site (Fig. 5D, Supplementary Fig. S7), providing the evidence of direct coupling between CDNF binding and nucleotide dynamics. Overall, our MD simulation data demonstrate that the interaction with C-CDNF is limiting the degree of freedom of BiP NBD lobes (Fig. 5B, Supplementary Fig. S6), which results in the stabilization of the ^NBD^K81-^NBD^E293 salt bridge (Fig. 5C), and subsequently preventing the nucleotide release from the active site (Fig. 5D).

### Neuronal survival assays demonstrate effect of CDNF mutations on human dopamine neuron rescue

To investigate whether pro-survival and neuroprotective properties of CDNF in human induced pluripotent stem cell (iPSC)-derived dopamine (DA) neurons rely on its interaction with BiP, we evaluated CDNF mutants (R117E, V118Y, A134Q, and E137A), which are deficient in BiP binding. Exposure of human DA neurons to 6-hydroxydopamine (6-OHDA) induced oxidative and ER stress followed by cell death. We found that WT CDNF in a dose-dependent manner protects significantly human DA neurons from 6-OHDA-induced cell death. In contrast, CDNF mutants deficient in BiP binding showed no significant protection against 6-OHDA-induced cell death, with survival rates similar to those observed in untreated neurons (Fig. 6). These findings suggest that R117E, V118Y, A134Q, and E137A mutations abolish CDNF neuroprotective activity, suggesting that BiP interaction might be important for neuroprotective activity of CDNF in human DA neurons.

**Figure 6.**
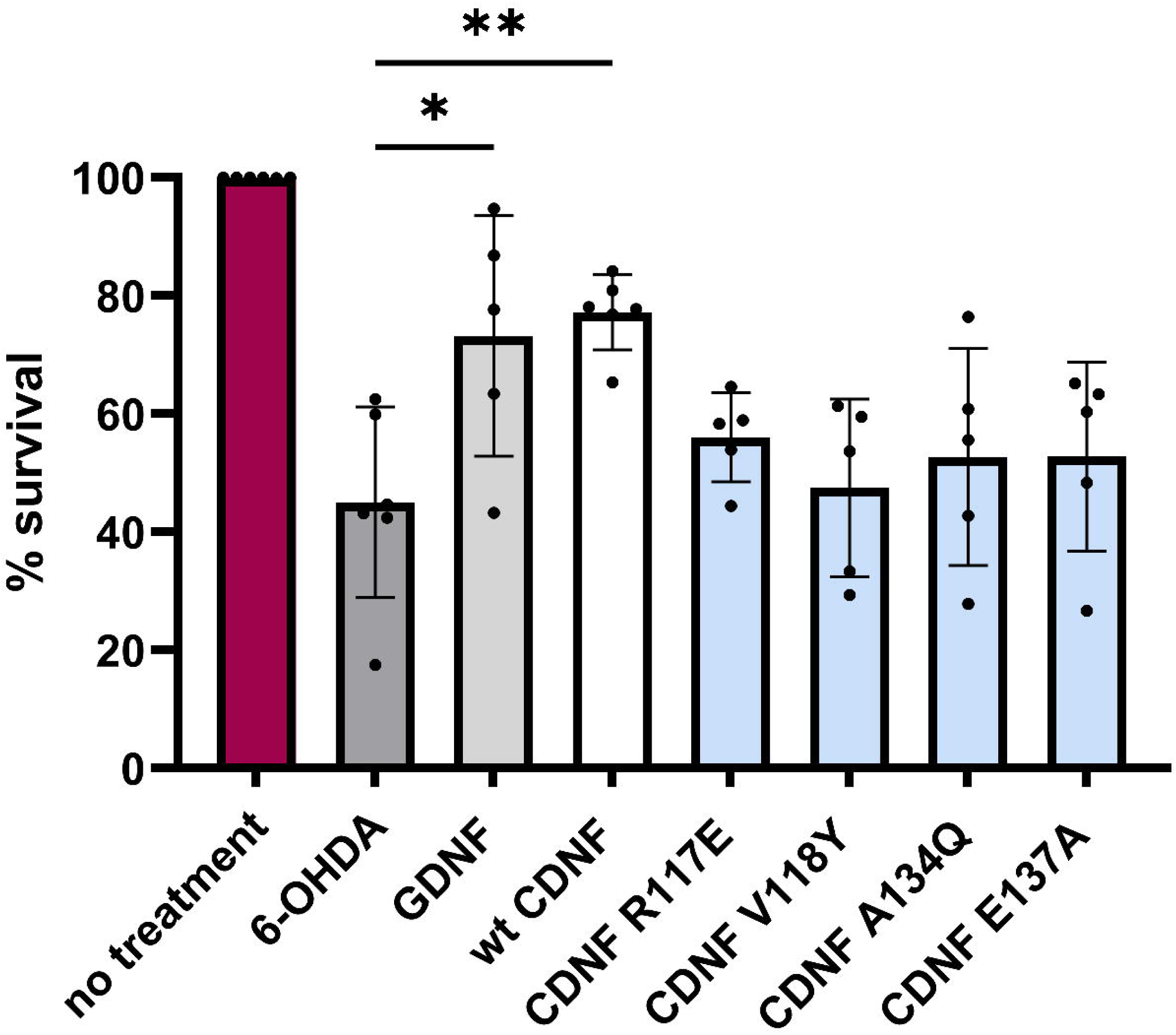
CDNF mutants deficient for BiP binding fail to protect human iPSC-derived DA neurons from 6-OHDA-induced cell death. Human iPSC-derived DA neurons were treated with 6-OHDA (50 µM) alone or in combination with either WT CDNF or BiP-binding deficient CDNF mutants (R117E, V118Y, A134Q, E137A) at 100 ng/ml. WT GDNF at concentration 10 ng/ml was used as positive control. Cell survival was measured as a percentage relative to untreated controls. WT CDNF displayed a significant dose-dependent protective effect, while BiP-binding deficient mutants did not show significant neuroprotection. Data are presented as mean ± SEM (n=4–5). Statistical significance was determined using one-way ANOVA with post hoc comparisons: ****p < 0.0001, **p < 0.01, *p < 0.05.

## 3. Discussion

CDNF is an ER-localized protein, interacting with ER stress-related proteins and UPR signaling regulators^7, 19, 22^. Although previous studies have identified CDNF as an ER stress-responsive protein with an *in vivo* neuroprotective role for DA neurons and motoneurons^1, 26^, the precise molecular mechanism of how CDNF regulates ER stress and protects neurons remains largely unclear. Here, we confirm the CDNF interaction with the main ER chaperone BiP through biochemical, biophysical, and structural studies, clarifying and extending the prior observations that CDNF can bind BiP^7^. We present the high-resolution crystal structures of CDNF in complex with BiP NBD. Our structural and functional studies clearly indicate that CDNF binds the ADP-bound state of BiP through its C-CDNF domain, but not the ATP-bound state of BiP, based on our MST binding measurements. This is in line with the highly different conformations of the ADP and ATP-bound states of BiP (Supplementary Fig. S2). We solved the structures of the CDNF-BiP complex with and without ADP, and in both cases Mg^2+^ and PO_4_^2-^ ions are bound. The ADP and PO_4_^2-^ bound structure describes the state after hydrolysis of ATP prior to release of the products, thus representing the natural state of the protein in its functional cycle, which CDNF likely interacts with in the cells, although there are no major conformational differences observed due to the presence or absence of ADP.

We show at atomic resolution that CDNF directly binds to BiP NBD, through charged hydrogen bonded interactions as well as through formation of hydrophobic pockets on the binding interface. Specifically, we identified the key interactions, some analogous to previously reported for MANF; mainly the CDNF R117 interaction with BiP D238 and CDNF E137 interactions with BiP N200 and T203. In addition, we show that CDNF V118 is packing against BiP V241, and that amino acid residues on the loop in C-CDNF containing the CXXC motif involving the C132-C135 disulphide bind to BiP through hydrophobic and hydrogen bonded interactions in the binding pocket. Thus, this CDNF loop (G129-E137) appears to have functional importance, and most likely the C132-C135 disulphide is important in stabilizing the loop conformation and thus the binding to BiP. The C132S and C135S mutants were shown to be defective in binding to BiP earlier in BiFC assays^22^. Hence, we note that there are several interactions involving the CDNF loop G129-E137 that were not accounted for in the earlier MANF-BiP complex analysis^21^ and that the CDNF binding interface does not only involve ionic hydrogen bonded interactions, but clearly also has a hydrophobic component. The importance of the G129-E137 loop is supported by our mutational studies, as mutation A134Q abolishes the interaction.

Recently, also SAXS-based model was reported suggesting that C-CDNF is responsible for binding to BiP^28^. However, the low-resolution model describes the binding mode of CDNF with BiP in a different orientation from ours. Only one CDNF residue identified in the SAXS data-based complex model, E137, is in common with the key interaction residues we have identified in our crystal structures, underscoring the fact that it is essential to have high-resolution data to be able to draw exact biological mechanistic conclusions. Given the importance of CDNF as a neurotrophic factor with the shown disease-modifying effect in PD and ALS models^25, 26^, it is important that the accurate interaction mechanisms of CDNF are identified with its target proteins. CDNF has been shown to interact with BiP^7^, as well as the UPR sensors PERK and IRE1α, which act as its intracellular receptors^22^, suggesting a dual role in the ER stress response.

We found the CDNF affinity to BiP to be in the micromolar range for the non-ATP bound states, with *K*_d_ of 1 μM by MST and 23.4 μM by SPR, the latter for the NBD. These values confirm a physiologically relevant affinity, which is abolished in the ATP-bound state (Fig. 1E). This nucleotide sensitivity can be directly connected with the effect of CDNF on BiP functional cycle and chaperone activity, where ATP binding induces an “open” conformation of SBD that weakens SBD substrate binding affinity, and due to conformational steric hindrance reduces the affinity for CDNF. The fact that we do not observe binding of CDNF in the presence of ATP suggests that ADP-bound state of BiP is required for the CDNF interaction, stabilizing this “closed” or high affinity substrate binding state. Our findings suggest that CDNF selectively engages BiP to enhance substrate-binding affinity by stabilizing the ADP-bound state.

Mutational and binding studies with purified proteins and in cell-based BiFC assays on the BiP-CDNF binding interface clearly confirm the importance of the interface residues. Moreover, the functional importance of the interaction was confirmed through nucleotide exchange and neuronal survival and neuroprotection assays.

We show with nucleotide exchange studies that CDNF acts as an antagonist of BiP nucleotide exchange, or nucleotide exchange inhibitor, affecting its chaperone cycle by slowing down the release of ADP, thus having the functional effect on BiP similar to MANF. We also demonstrate that N-CDNF does not contribute to BiP binding based on pull down experiments, nor does it have an effect on the nucleotide exchange, unlike the C-CDNF or the full-length CDNF. Thus, the main BiP regulatory domain of CDNF clearly is C-CDNF. We also show CDNF can counteract the nucleotide exchange factor Grp170 action and that mutations of CDNF at the BiP binding site reduce its activity as a nucleotide exchange inhibitor. The effect of CDNF on Grp170 activity is very clear (Fig. 4D), and we propose that the main function of CDNF is not only to stabilize BiP in ADP-bound conformation but also to counteract the Grp170 and other NEF functions at the same time possibly by restricting the NBD conformational flexibility (Supplementary Fig. S8). Furthermore, CDNF and MANF also bind to the same site as is found for the J-domain-Hsp70 interaction, which is required for substrate delivery and acceleration for ATP hydrolysis^33^. In fact, the BiP residue R197 interacting with CDNF (Fig. 2B) has been implicated as key residue for BiP-J-domain interaction^34^. It is an intriguing possibility that CDNF (and MANF) may also counteract the J-domain function, which remains to be studied further.

The effect of CDNF on the conformational dynamics of BiP NBD is supported further by our MD simulation studies, which suggested reduced conformational flexibility of BiP NBD and inhibition of ADP release by CDNF. Based on the MD results, we show the effect of CDNF on nucleotide release to be directly related at the molecular level to the reduced conformational flexibility of the NBD. This is also expected in the presence of NEFs, which can induce large conformational changes to BiP NBD^35, 36^ (Supplementary Fig. S8). Moreover, our mechanistic insights suggest that restricted mobility of the BiP NBD by CDNF stabilizes the highly conserved inter-lobe salt bridge (K81-E293), which was previously reported to play a crucial role in nucleotide exchange in eucaryotic BiP NBD^37, 38^, thereby lowering the likelihood of ADP release from the binding site.

Importantly, we here also show the effect of CDNF mutations on the CDNF ability to protect and rescue human stem cell-derived dopamine neurons in a chemically induced cellular stress and neurodegeneration condition. Drawing together this and our other data^22^, we propose a model for CDNF function with roles in BiP chaperone cycle regulation as well as in the regulation of UPR, both of which can promote neuronal cell survival, through direct interactions with both BiP and UPR sensor molecules PERK and IRE1α, exact mechanism of which remains to be confirmed in more detail. Previous studies with mouse and human cultured DA neuronal models demonstrated that both PERK-binding deficient CDNF mutant (K122A) and IRE1α-binding deficient CDNF mutants (Y58F, Y102F) are functional in BiP binding but lacked neuroprotective activity upon 6-OHDA treatment^22^. In contrast, cysteine loop mutants (C132S, C135S) retained neuroprotective efficacy in tunicamycin-induced ER stress and 6- OHDA models comparable to WT CDNF and GDNF but showed no binding to BiP. These data suggest that both BiP interaction and UPR receptor interactions have importance in mediating protective functions of CDNF in human DA neurons. It should be noted, however, that some CDNF mutants lacking the ability to bind BiP retain full neuroprotective activity.

## 4. Methods

### Plasmid construction

The following constructs were made for the *E. coli* expression system. Human BiP NBD (aa 26-405) gene was cloned into pHYRSF53^39^ (a kind gift from Dr. Hideo Iwai) between BamHI and HindIII sites, which results in expressing N-terminally 6×His- and Smt3-tagged BiP NBD (6His-Smt3-BiP NBD). Human full-length BiP (BiP FL, aa 25-654), BiP NBD (aa 25-407), and BiP SBD (aa 418-654) genes were separately cloned into pHAT3^40^ between SpeI and HindIII sites, which results in expressing N-terminally 6×His-tagged BiP proteins with a human rhinovirus 3C protease cleavage site between them (6His-3C-BiP FL, NBD, and SBD). Full-length CDNF (FL-CDNF, aa 1-161), N-CDNF (aa 1-103), and C-CDNF (aa 101-161) genes were also separately cloned into pHAT3 in the same way to express 6His-3C-CDNF proteins. CDNF mutant constructs (R117E, V118Y, A134Q, and E137A) were made by mutagenesis using the plasmid pHAT3 containing the FL-CDNF gene as a template. A plasmid pCA833 was kindly provided by Dr. Claes Andreasson (Stockholm University) for expressing 6His-Smt3-Grp170 (aa 33-999). This plasmid is a derivative of the pSUMO-YHRC vector, which was described previously^41^. All the constructs were validated by Sanger sequencing.

The construction of the plasmids for the BiFC system (pEZY BiFC Jun-NV, pEZY BiFC Max- CV, pEZY BiFC Fos-CV, pEZY BiFC Grp78-NV(CV), pEZY BiFC pre-CDNF-NV(CV), pEZY BiFC pre-PERK-NV(CV), and pEZY BiFC IRE1α -NV(CV)) was previously described in detail^20, 22, 42^. Mutant versions of CDNF (R117E, V118Y, A134Q, Q137A, and V118Y+E137A) were generated using site-directed inverse PCR mutagenesis with the pEZY BiFC CDNF-NV(CV) plasmid as a template. Inverse PCR was performed using Phusion High-Fidelity DNA Polymerase (Thermo Fisher Scientific) with mutagenic primers designed using the PrimerX tool (https://www.bioinformatics.org/primerx/). Following PCR amplification, the template plasmid was digested with DpnI (New England Biolabs) at 37°C for 2 hours to eliminate parental DNA. The digested products were purified using a PCR cleanup kit (Qiagen) and transformed into DH5α *E. coli* cells via heat shock at 42°C for 45 s. Cells were plated on ampicillin (100 μg/ml) LB-agar plates and incubated overnight at 37°C. Colonies were screened via colony PCR and successful clones were confirmed by Sanger sequencing (Eurofins Genomics). The V118Y-E137A double mutant was generated using the V118Y construct as a template and subjected to the same mutagenesis protocol.

For recombinant protein production with the mammalian CHO cell expression, the pre-SH-CDNF coding sequence, containing an N-terminal signal peptide (pre) and twin Strep-II + HA (SH) tag, was cloned into the pcDNA5/FRT/TO vector (Invitrogen) to generate the pcDNA5/FRT/TO pre-SH-CDNF construct, as previously described^7^. Mutations R117E, V118Y, A134Q, Q137A, and V118Y-E137A were introduced into the pcDNA5/FRT/TO pre-SH-CDNF plasmid using site-directed inverse PCR mutagenesis as described above. The confirmed mutant plasmids were used for transient transfection in cells for protein production.

### Protein expression and purification from mammalian cells

For initial binding interaction studies, WT CDNF and its mutant variants (R117E, V118Y, A134Q, Q137A, and V118Y+E137A) were expressed in a mammalian system using a CHO-derived cell line, as described before^42, 43^. The recombinant proteins were produced by Icosagen (https://www.icosagen.com/) using an optimized transient expression platform. Additionally, some CDNF proteins were produced by Biovian (Turku, Finland) following a similar methodology. In brief, codon-optimized CDNF cDNAs were cloned into pQMCF-T expression vectors and transiently transfected into CHO-derived protein production cells. Following expression, proteins were isolated from the culture medium through a sequential purification process involving a 5 ml Q FF column, followed by a 1 ml SP HP column. The buffer was exchanged to PBS (pH 7.4) via size-exclusion chromatography. Protein purity was assessed using SDS-PAGE with Coomassie staining and validated through immunoblotting with a polyclonal rabbit anti-CDNF antibody (300-100, Icosagen).

### Protein expression and purification from *E. coli*

All the BiP constructs for the *E. coli* expression system were expressed in the Rosetta (DE3) *E. coli* strain (Merck Millipore). All the cultures except for the 6His-3C-BiP FL construct were grown in LB medium containing 50 µg/ml kanamycin for the 6His-Smt3-BiP NBD construct or 100 µg/ml ampicillin for the other constructs at 37°C until OD_600nm_ reached 0.8, and the proteins were expressed at 18°C for 20 h after adding 0.2-0.5 mM IPTG for induction. For the 6His-3C-BiP FL construct, TB medium was used instead of LB medium, and the culture was grown until OD_600nm_ reached 1.8 before IPTG induction with the same condition. The cells were harvested by centrifugation, and the pellets were stored at -20°C until further use.

All the CDNF constructs for the *E. coli* expression system were expressed in the Rosetta-gami 2(DE3) *E. coli* strain (Merck Millipore). Cultures were grown in LB medium containing 100 µg/ml ampicillin, 25 µg/ml chloramphenicol, and 10 µg/ml tetracycline at 37°C until OD_600nm_ reached 0.6, and the proteins were expressed at 30°C for 18 h after adding 0.2 mM IPTG for induction. The cells were harvested by centrifugation, and the pellets were stored at -20°C until further use.

The 6His-Smt3-Grp170 construct was expressed in the Rosetta (DE3) *E. coli* strain (Merck Millipore). It was grown in LB medium containing 50 µg/ml kanamycin at 37°C until OD_600nm_ reached 0.5, and the proteins were expressed at 18°C for 20 h after adding 0.5 mM IPTG for induction. The cells were harvested by centrifugation, and the pellets were stored at -20°C until further use.

Cell pellets of the 6His-Smt3-BiP NBD construct were lysed by sonication in resuspension buffer A (50 mM Tris-HCl, pH 8.0, 500 mM NaCl, 20 mM imidazole). The lysate was centrifuged at 20,000 × g for 30 min, and then the supernatant was incubated with HisPur Ni-NTA Resin (ThermoFisher) for 60 min at 4°C. After washing the resin with wash buffer A (50 mM Tris-HCl, pH 8.0, 150 mM NaCl, 20 mM imidazole), the protein was eluted with elution buffer A (50 mM Tris-HCl pH 8.0, 150 mM NaCl, 400 mM imidazole). The buffer of the protein solution was exchanged to wash buffer A by concentrating the solution with 10 kDa NMWL Amicon Ultra centrifugal filters and diluting it with dilution buffer A (50 mM Tris-HCl, pH 8.0, 150 mM NaCl). The 6His-Smt3 tag was cleaved by incubating the sample with the homemade 6His-Ulp1 protein and 1 mM DTT overnight at 4°C. The sample was then incubated again with HisPur Ni-NTA Resin (ThermoFisher) for 60 min at 4°C, and the unbound fraction was collected. The sample was further purified with a Superdex 75 10/300 GL column (GE Healthcare) equilibrated with running buffer A (10 mM Tris-HCl, pH 7.5, 100 mM NaCl, 5 mM MgCl_2_), concentrated, flash-frozen, and stored at -80°C.

Cell pellets of the 6His-3C-BiP FL, NBD, and SBD constructs were lysed by sonication in resuspension buffer B (50 mM Tris-HCl, pH 8.0, 500 mM NaCl, 10 mM imidazole). The lysate was centrifuged at 20,000 × g for 30 min, and then the supernatant was incubated with HisPur Ni-NTA Resin (ThermoFisher) for 60 min at 4°C. After washing the resin with wash buffer A, the protein was eluted with elution buffer A. The sample was further purified with a Superdex 75 10/300 GL column (GE Healthcare) for the NBD and SBD constructs and with a Superdex 200 10/300 GL column (GE Healthcare) for the FL construct equilibrated with running buffer B (10 mM HEPES, pH 7.4, 150 mM NaCl), respectively, then concentrated, flash-frozen, and stored at -80°C.

Cell pellets of the 6His-3C-CDNF constructs (FL-, N-, and C-CDNF and FL-CDNF mutants) were lysed by sonication in resuspension buffer C (50 mM Tris-HCl, pH 8.0, 500 mM NaCl, 20 mM imidazole, 1 mM PMSF). The lysate was centrifuged at 20,000 × g for 30 min, and then the supernatant was incubated with HisPur Ni-NTA Resin (ThermoFisher) for 60 min at 4°C. After washing the resin with wash buffer B (50 mM Tris-HCl, pH 8.0, 500 mM NaCl, 20 mM imidazole), the protein was eluted with elution buffer A. The buffer of the protein solution was exchanged to wash buffer A by concentrating the solution with 3 kDa NMWL Amicon Ultra centrifugal filters and diluting it with dilution buffer A. The tag was cleaved by incubating the sample with the in-house made GST-tagged human rhinovirus 3C protease protein and 0.1 mM DTT overnight at 4°C. The sample was then purified with a Superdex 75 10/300 GL column (GE Healthcare) equilibrated with running buffer C (10 mM HEPES, pH 7.4, 100 mM NaCl), concentrated, flash-frozen, and stored at -80°C.

Cell pellets of the 6His-Smt3-Grp170 construct were lysed by sonication in resuspension buffer D (50 mM Tris-HCl, pH 8.0, 500 mM NaCl, 20 mM imidazole, 2 mM MgCl_2_, 0.1 mM TCEP, 10% glycerol, 1 mM PMSF, 0.1 mg/ml DNase I). The lysate was centrifuged at 20,000 × g for 30 min, and then the supernatant was incubated with HisPur Ni-NTA Resin (Thermo Fisher) for 60 min at 4°C. After sequentially washing the resin with wash buffer C (50 mM Tris-HCl, pH 8.0, 500 mM NaCl, 20 mM imidazole, 2 mM MgCl_2_, 0.1 mM TCEP, 10% glycerol), wash buffer D (25 mM Tris-HCl, pH 8.0, 150 mM NaCl, 10 mM imidazole, 0.1 mM TCEP), wash buffer E (25 mM Tris-HCl, pH 8.0, 1000 mM NaCl, 10 mM imidazole, 0.1 mM TCEP), wash buffer F (25 mM Tris-HCl, pH 8.0, 150 mM NaCl, 10 mM imidazole, 0.1 mM TCEP, 10 mM MgCl_2_, 3 mM ATP, 0.01 mg/ml RNase A), wash buffer G (500 mM Tris-HCl, pH 8.0, 150 mM NaCl, 10 mM imidazole, 0.1 mM TCEP), wash buffer E, and wash buffer D, the protein was eluted with elution buffer B (25 mM Tris-HCl, pH 8.0, 150 mM NaCl, 400 mM imidazole, 1 mM DTT). The buffer of the protein solution was exchanged to decrease the imidazole concentration to 20 mM by concentrating the solution with 10 kDa NMWL Amicon Ultra centrifugal filters and diluting it with dilution buffer B (10 mM Tris-HCl, pH 8.0, 150 mM NaCl, 1 mM DTT). The 6His-Smt3 tag was cleaved by incubating the sample with the in-house made 6His-Ulp1 protein overnight at 4°C. The sample was then incubated again with HisPur Ni-NTA Resin (ThermoFisher) for 60 min at 4°C, and the unbound fraction was collected. The sample was further purified with a Superdex 200 10/300 GL column (GE Healthcare) equilibrated with running buffer D (10 mM HEPES, pH 7.4, 150 mM KCl, 0.2 mM TCEP), concentrated, flash-frozen, and stored at -80°C. Mass spectrometry analysis of CDNF purified from *E. coli* confirmed the molecular weight corresponding to reduced cysteines, and the protein was soluble and functional.

### C-CDNF peptide production

The 61 amino acid C-CDNF peptide was also produced and purified by Shanghai peptide Co. Ltd., Shanghai, China. The purity was analyzed by HPLC and the peptide structure verified by mass-spectrometry and NMR spectroscopy. The biological activity of the C-CDNF was confirmed using microinjection to mouse SCG sympathetic neurons and testing their neuronal survival-promoting activity^44^. The comparison of survival-promoting activity of C-CDNF to the effect of full-length CDNF and nerve growth factor (NGF) showed that C-CDNF exerts full biological activity.

### Pull-down assays

To test binding of CDNF to different BiP domains, 25 µM of 6His-3C-BiP FL, 6His-3C-BiP NBD, and 6His-3C-BiP SBD in assay buffer A (10 mM Tris-HCl, pH 7.4, 100 mM NaCl, 2 mM MgCl_2_, 40 mM imidazole) were separately incubated with Ni-NTA agarose (QIAGEN) for 1 h at room temperature. For a control, just assay buffer A was incubated instead. After washing the resin four times with assay buffer A, 100 µM of CDNF purified from CHO cells in assay buffer A was incubated with the resin for 1 h at room temperature. The resin was again washed four times with assay buffer A, and the bound proteins were eluted with elution buffer A. The eluted proteins were analyzed by SDS-PAGE with 4–20% Mini-PROTEAN TGX Stain-Free protein gels (Bio-Rad) and with the stain-free detection protocol using the Gel Doc EZ imager (Bio-Rad).

To test binding of different domains of CDNF to BiP NBD, 25 µM of 6His-3C-BiP NBD in assay buffer B (10 mM Tris-HCl, pH 7.4, 100 mM NaCl, 10 mM MgCl_2_, 20 mM imidazole) was incubated with some aliquots of Ni-NTA agarose (QIAGEN) for 1 h at room temperature. For controls, just assay buffer B was incubated instead. After washing the resin four times with assay buffer B, 100 µM of N-CDNF and C-CDNF purified from the *E. coli* system in assay buffer B were separately incubated with each aliquot of the resin for 1 h at room temperature. The resin was again washed four times with assay buffer B, and the bound proteins were eluted with elution buffer A. The eluted proteins were run through 4–20% Mini-PROTEAN TGX Stain-Free protein gels (Bio-Rad) by SDS-PAGE, stained with InstantBlue Coomassie Protein Stain (Abcam), and imaged with iBright FL1500 (ThermoFisher).

To test binding of different mutants of CDNF to BiP NBD, 25 µM of 6His-3C-BiP NBD in assay buffer B was incubated with some aliquots of Ni-NTA agarose (QIAGEN) for 1 h at room temperature. For controls, just assay buffer B was incubated instead. After washing the resin four times with assay buffer B, 100 µM of wild-type CDNF and the mutants purified from CHO cells in assay buffer B were separately incubated with each aliquot of the resin for 1 h at room temperature. The resin was again washed four times with assay buffer B, and the bound proteins were eluted with elution buffer A. The eluted proteins were analyzed by SDS-PAGE with 4–20% Mini-PROTEAN TGX Stain-Free protein gels (Bio-Rad) and with the stain-free detection protocol using the Gel Doc EZ imager (Bio-Rad).

### Analytical size-exclusion chromatography

To see the complex formation of BiP NBD and CDNF in solution, BiP NBD and CDNF purified from CHO cells were mixed and incubated at 100 µM for each for 30 min at room temperature in running buffer E (PBS, pH 7.4, 2 mM MgCl_2_). For controls, only BiP NBD and only CDNF (CHO) both at 100 µM were also incubated with the same condition. The samples were then run through a Superdex 200 Increase 3.2/300 column (Cytiva) connected to the ÄKTA pure chromatography system (Cytiva) equilibrated with running buffer E at 4°C. The eluted fractions were analyzed by SDS-PAGE with 4–20% Mini-PROTEAN TGX Stain-Free protein gels (Bio-Rad) and with the stain-free detection protocol using the Gel Doc EZ imager (Bio-Rad).

### Surface plasmon resonance

Surface plasmon resonance (SPR) analysis was conducted with Biacore T100 or T200 systems (Cytiva) for measuring affinity between BiP NBD and WT CDNF and for comparing affinity between BiP NBD and different CDNF mutants all purified from the *E. coli*, with running buffer F (20 mM HEPES, pH 7.4, 150 mM NaCl, 2 mM MgCl_2_, 0.02% Tween 20). BiP NBD was immobilized on a CM5 chip (Cytiva) as a ligand by amine coupling following the standard procedure provided by the manufacturer, and target response was set to 5000 response units (RU). Then serial dilutions of CDNF (for WT CDNF affinity measurement) or 50 µM of different CDNF constructs (WT and the mutants for response level comparison) were sequentially injected as analytes with the flow rate of 30 µl/min for a contact time of 90 s followed by 300 s dissociation time. After each injection, the surface was regenerated by running 1 M NaCl for a contact time of 30 s with the same flow rate. Steady-state binding levels were used for both calculating the dissociation constant (*K*_d_) of BiP NBD-CDNF interaction and for comparing the BiP binding to different CDNF mutants.

### Microscale thermophoresis

Binding affinities of recombinant proteins were measured at 25°C using the Monolith NT.115 instrument (NanoTemper Technologies GmbH, Germany). The recombinant proteins used in MST experiments were purified from CHO cells, while BiP was obtained from *E. coli*. BiP-His protein was labeled using the His-Tag Labeling Kit RED-tris-NTA (MO-L008), and CDNF, BiP NBD proteins with the amine-reactive Monolith Protein Labeling Kit RED-NHS 2nd Generation (L011). Unbound dye was removed after labeling using Zeba Spin Desalting Columns (89882, Thermo Fisher Scientific) following the manufacturer’s instructions. For all assays, the proteins were maintained at a concentration of 20 nM in MST buffer, consisting of 10 mM sodium phosphate (pH 7.4), 1 mM MgCl_2_, 3 mM KCl, 150 mM NaCl, and 0.05% Tween-20. Measurements were performed using premium-coated capillaries (NanoTemper Technologies GmbH), with an optimized LED power setting of 100% for fluorescence detection. Binding curves were generated using 12-14 concentration points, with each point representing the average of 3-5 independent experiments. The affinity curves for distinct CDNF mutants (each measured 3-5 in a row) were produced by MO.Affinity Analysis software (NanoTemper Technologies). Data analysis involved determining normalized fluorescence changes (ΔFnorm), calculating standard deviations, and estimating dissociation constants (Kd) along with error margins. The data was analyzed with MO.Affinity Analysis software (versions 2.3 and 2.2.4) and GraphPad Prism (versions 8.0.2 and 9).

### Crystallization

C-CDNF produced by Shanghai peptide Co. Ltd. that had been dissolved in PBS was dialyzed against 10 mM Tris-HCl, pH 7.5, 100 mM NaCl, 5 mM MgCl_2_. BiP NBD and C-CDNF were mixed at 1:1.3 molar ratio (final 360 and 470 µM, respectively) with or without 2 mM ADP and used for crystallization screening by the sitting drop method with 100 nl of protein solution and 100 nl of well solution at 20°C. Thin plate crystals for the BiP NBD and C-CDNF complex without ADP were obtained from a drop containing a well solution composed of 100 mM HEPES, pH 7.5, 200 mM NaCl, 25% (w/v) PEG3350. Thin plate crystals for the complex with ADP were obtained with a well solution composed of 100 mM Bis-Tris, pH 5.5, 100 mM NaCl, 25% (w/v) PEG3350. The crystals were briefly soaked into the well solution containing 15% (v/v) glycerol with or without 2 mM ADP for the complex with or without ADP, respectively, and immediately stored in liquid nitrogen.

### Data collection and structure determination

Diffraction data of the complex of BiP NBD and C-CDNF without ADP were collected at the I24 beamline in the Diamond Light Source synchrotron and processed by the autoPROC pipeline^45^, which includes XDS^46^ for indexing and integration, POINTLESS^47^ for space group determination, and AIMLESS^48^ for scaling and merging. The structure was solved initially by molecular replacement using Phaser^49^ with chain A (protein residues only) of PDB 5F0X^32^, a known BiP NBD structure, as a search model. Then the C-CDNF model was built using ARP/wARP^50^ into the solved BiP NBD structure. Further refinement and manual model building were conducted iteratively with REFMAC5^51^ and Coot^52^, respectively. MolProbity^53^ was used for structure validation.

Diffraction data of the complex of BiP NBD and C-CDNF with ADP were collected at the ID30A-3 beamline in European Synchrotron Radiation Facility (ESRF) and processed by the autoPROC pipeline. The output data from XDS were reprocessed with XSCALE for scaling and to merge Friedel pairs and then merged with AIMLESS. The structure was solved initially by molecular replacement using MOLREP^54^ with the BiP NBD and C-CDNF complex without ADP as a search model. Further refinement and manual model building were conducted iteratively with REFMAC5 and Coot, respectively. MolProbity was used for structure validation.

### Bimolecular fluorescence complementation assay (BiFC)

HEK293 cells were plated on poly-D-lysine-coated coverslips (P0899, Sigma-Aldrich) and co-transfected 48 hours later with pEZY BiFC N-Venus and C-Venus plasmids, using jetPEI transfection reagent (101, Polyplus-transfection), according to the manufacturer’s guidelines. Following transfection, the cells were fixed with 4% paraformaldehyde (PFA), washed with PBS, and permeabilized using 0.1% Triton X-100 in PBS (PBS-T). Nuclei and endoplasmic reticulum (ER) were stained using the ER-ID Red assay kit (ENZ-51026-K500, Enzo Life Sciences), which includes Hoechst 33342 for nuclear staining and ER-ID Red for ER visualization. Coverslips were mounted with ProLong Diamond Antifade Mountant (P36965, Thermo Fisher Scientific), and images were acquired on a Leica SP8 STED confocal microscope, equipped with a 96x glycerol immersion objective. Image acquisition was managed with Leica Application Suite X (LASX) software, and post-processing (including uniform brightness and contrast adjustments) was carried out using CorelDRAW 2018.

### Fluorescence-activated cell sorting

HEK293 cells were seeded at a density of 0.1 to 0.2 × 10^6^ cells per well in 12-well plates. The following day, cells were transfected with the BiFC NV and BiFC CV constructs for the proteins of interest. After 48 hours of transfection, cells were dissociated using TrypLE without phenol red (12604021, Thermo Fisher) and neutralized with media (Opti-Mem without phenol red + 10% FBS). The cell suspensions were centrifuged, the supernatant discarded, and the pellet resuspended in PBS. The cells were then filtered through a cell strainer cap and collected in polypropylene tubes (both from BD Biosciences) to remove clumps. Fluorescence was analyzed using a FACS Fortessa flow cytometer (BD Biosciences).

### Nucleotide exchange assay

Effects of CDNF on the release of a fluorescent ADP analog MABA-ADP (NU-893-MNT, Jena Bioscience) from BiP were tested essentially as has been published before^21^. Briefly, 5 µM BiP NBD or 6His-3C-BiP FL was mixed with 5 µM MABA-ADP in equal volume in assay buffer B (50 mM HEPES-KOH, pH 7.4, 100 mM NaCl, 10 mM MgCl_2_) and incubated in the dark for 90 min at 30 °C for the BiP NBD sample and for 20 h at room temperature for the BiP FL sample, respectively. The BiP-MABA-ADP solution was mixed in a 100 µl quartz cuvette (Starna Scientific) with the equal volume of nucleotide exchange solution (250 µM ATP and 200 µM CDNF variant as indicated, all from the *E. coli* expression system, in assay buffer B), and the release of MABA-ADP was immediately monitored by the decrease in MABA-ADP fluorescence at excitation 360 nm and emission 440 nm using Cary Eclipse Spectrophotometer (Agilent Technologies). Final concentrations of reaction components: BiP 1.25 μM, MABA-ADP 1.25 μM, ATP 125 μM, and 100 µM CDNF variant as indicated. For the experiments with Grp170 and Ca^2+^, 1 mM CaCl_2_ was included in the BiP-MABA-ADP solution before the 90 min incubation, and 2.5 µM Grp170 was included in the nucleotide exchange solution. Thus, the final concentrations of Ca^2+^ and Grp170 were 0.5 mM and 1.25 µM, respectively. The curves were fitted to the one phase exponential decay function in Prism 10 (GraphPad), and the MABA-ADP release rates (*k*_off_) were determined from three to four independent experiments.

### Molecular dynamics simulations

Large-scale atomistic molecular dynamics (MD) simulations were conducted based on the high-resolution crystal structure of human BiP nucleotide binding domain bound by the C-terminal of CDNF. To assess the impact of C-CDNF binding on the conformational dynamics of BiP NBD, we constructed an additional simulation setup by removing C-CDNF from the “full” model system. Furthermore, to simulate the ADP release from BiP NBD, we created additional setups from the ADP-bound crystal structure of BiP NBD both in the presence and absence of C-CDNF (latter PDBID: 5EVZ). Each system was solvated in a cubic water box using the TIP3P water model^55^ with box dimensions of 107×107×107 (Å^3^) and physiological salt concentration of 150 mM NaCl, yielding a total system size of approximately 120,000 atoms.

CHARMM36 force field^56^ was applied in our simulations, where all model systems adopted the identical simulation protocol. At the first step, we performed initial minimization with harmonic restraints on protein heavy atoms and, in ADP-bound model setups, on ADP molecules (force constant k=4.2*10^4^kJ/(mol*nm^2^). The minimization was performed for 1000 steps using conjugate gradient algorithm, as implemented in NAMD software package (v2.14)^57^. This was followed by subsequent minimization in GROMACS (v.2022.3)^58^ using steepest-descent algorithm with the force tolerance of 250 kJ/(mol*nm), imposing the same restraints with a reduced force constant of 2.0*10^4^ kJ/(mol*nm^2^). Subsequently, two sequential equilibration stages were carried out in NPT ensemble: a 10-ns run with restraints on the same atoms, followed by a 10-ns equilibration with restraints applied only to the C_α_ atoms of the protein backbone. Temperature control (T=310 K) during equilibration was achieved using the V-rescale thermostat^59^, while constant pressure was maintained at 1 atm via C-rescale barostat^60^. The production phase included unrestrained MD run for 1 μs with Nose-Hoover thermostat and Parrinello-Rahman barostat^61, 62^. Each of the model systems was simulated with 8 replicates, where replicas 2 through 8 were generated by extending of the first equilibration phase by 1, 3, 5, 6, 7, 8, and 9 ns, respectively. The total simulation time in the present work accounted for ∼32 microseconds. Non-bonded (Wan der Vaals) interactions were treated via Verlet cutoff scheme^63^, while long-range electrostatic interactions were accounted for using the particle mesh Ewald approach^64^. The trajectories were sampled using the “leapfrog” integration scheme with a 2-fs timestep. All covalent bonds to hydrogen atoms were restrained with the LINCS algorithm^65^. The trajectories were analysed with VMD analysis software^66^ and MDAnalysis Python package^67^. The figures of molecular structures were created with VMD and PyMOL (Schrödinger LCC) visualization tools.

Analysis of distances (Fig. 5A, B, D) and molecular contacts between NBD and C-CDNF (Supplementary Fig. S4) was performed from combining all simulation replicas into a single 8 μs dataset. The pair of residues was considered to be in ”contact” if any pair of atoms in these residues approach each other for less than 5 Å. The dynamical behavior of ADP in the binding pocket was evaluated as the distance between the ADP purine ring (atoms N1, C2, N3, C4, C5, C6) and the C *α* carbon of ^NBD^Ser365 (see Fig. 5D, right panel). The principal component analysis (PCA) was performed using GROMACS “anaeig” tool on a reduced dataset where we neglected first 300 ns from each replica. The Supplementary movies S1 and S2 were obtained from determining the smallest and largest displacement of protein C *α* carbon atoms along the first principal component, and interpolating between these two extreme values, as implemented in GROMACS “anaeig” tool. Porcupine plots (Fig. 5B) were generated using the PorcupinePlot (v1.0) plugin for VMD, based on C_α_ atom displacements between structures with the smallest and largest projections along the first principal component.

### Human iPS-cell derived dopamine neurons and treatment with CDNF and its mutants

iCell® DopaNeurons (R1088, FUJIFILM Cellular Dynamics) were seeded according to FUJIFILM Cellular Dynamics user protocol onto the 96-well plates coated with poly-L-ornithine (P3655, Sigma-Aldrich) and laminin (L2020, Sigma-Aldrich). Equal volumes of cell suspension were plated onto the center of the dish. The cells were grown for 5 days in iCell Neural Base Medium 1+ iCell Neural Supplement B (M1010, M1029, FUJIFILM Cellular Dynamics). Then the cells were treated with 6-hydroxydopamine hydrochloride (6-OHDA, 50 µM, H4381, Sigma Aldrich) and recombinant protein CDNF mutants (100 ng/ml), without any trophic factor or with GDNF (10 ng/ml) (P-103-100, Icosagen), as negative and positive control, respectively. After three days, the cells were fixed and stained with anti-tyrosine hydroxylase (TH) antibody (MAB318, Millipore Bioscience Research Reagents). Images were acquired by ImageXpress Nano automated imaging system. Immunopositive neurons were counted by CellProfiler software, and the data was analyzed by CellProfiler analyst software. The results are expressed as % of cell survival compared non-treated neurons.

## Acknowledgements

MS and TK acknowledge funding for the project from Jane and Aatos Erkko Foundation and MS also funding from the Research Council of Finland (grant No 343299). OZ acknowledges funding from the Research Council of Finland (grant No 362411). OS, MS and TK are grateful to Dr. Vera Kovaleva for help and stimulating discussions. OZ thanks Prof. Vivek Sharma and Dr. Outi Haapanen for fruitful discussions. The facilities and expertise of the HiLIFE Crystallization unit at the University of Helsinki, a member of FINStruct and Biocenter Finland are gratefully acknowledged. We acknowledge CSC IT Center for Science (Finland), and the IT Support from the University of Helsinki for computational resources. Coordinates for the BiP-CDNF structures and structure factors have been deposited to the Protein Data Bank with accession codes 9H0C and 9I5Y.

