## Supplementary materials for "Structure and functional analysis of CDNF-BiP interaction reveal a role in endoplasmic reticulum proteostasis regulation"

**Supplementary material. Fudo et al.**

**Figure S1.** Surface plasmon resonance assay on CDNF binding to BiP NBD. A) sensogram of CNDF half-dilution concentration series as indicated in the figure B) Binding curve from concentration series for *K*_d_ determination.

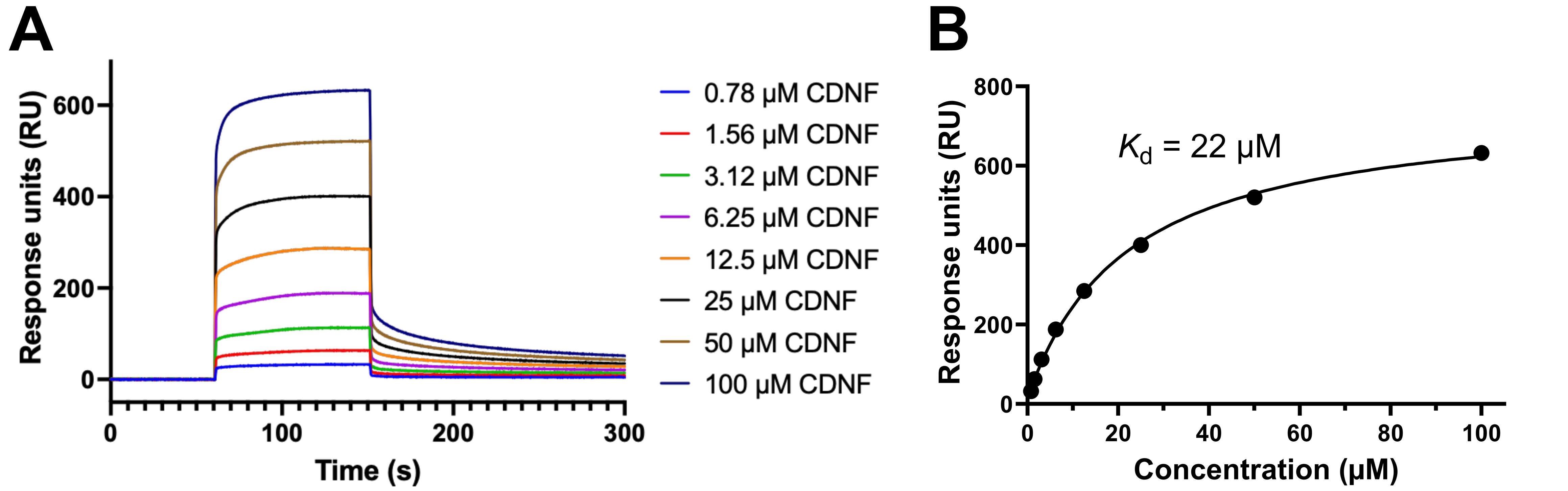

**Figure S2. Structural alignment of CNDF-BiP complex with full length BiP apo and ATP bound forms.** A) CNDF complex aligned with apo-BiP with SBD (red, linker; yellow, SBD) (PDB 6HAB), other colours as in Figure 2. (e.g. green, C-CDNF). B) Alignment with ATP-bound full length BiP (PDB: 5E84), yellow SBD, including the C-terminal α-helical “lid” subdomain (ATP bound structure in “docked” or “open” conformation). Note the much better alignment with the NBD domain in apo /ADP bound conformation of BiP. Bound C-CDNF clashes with the position of the linker between NBD and SBD in ATP bound conformation of BiP.

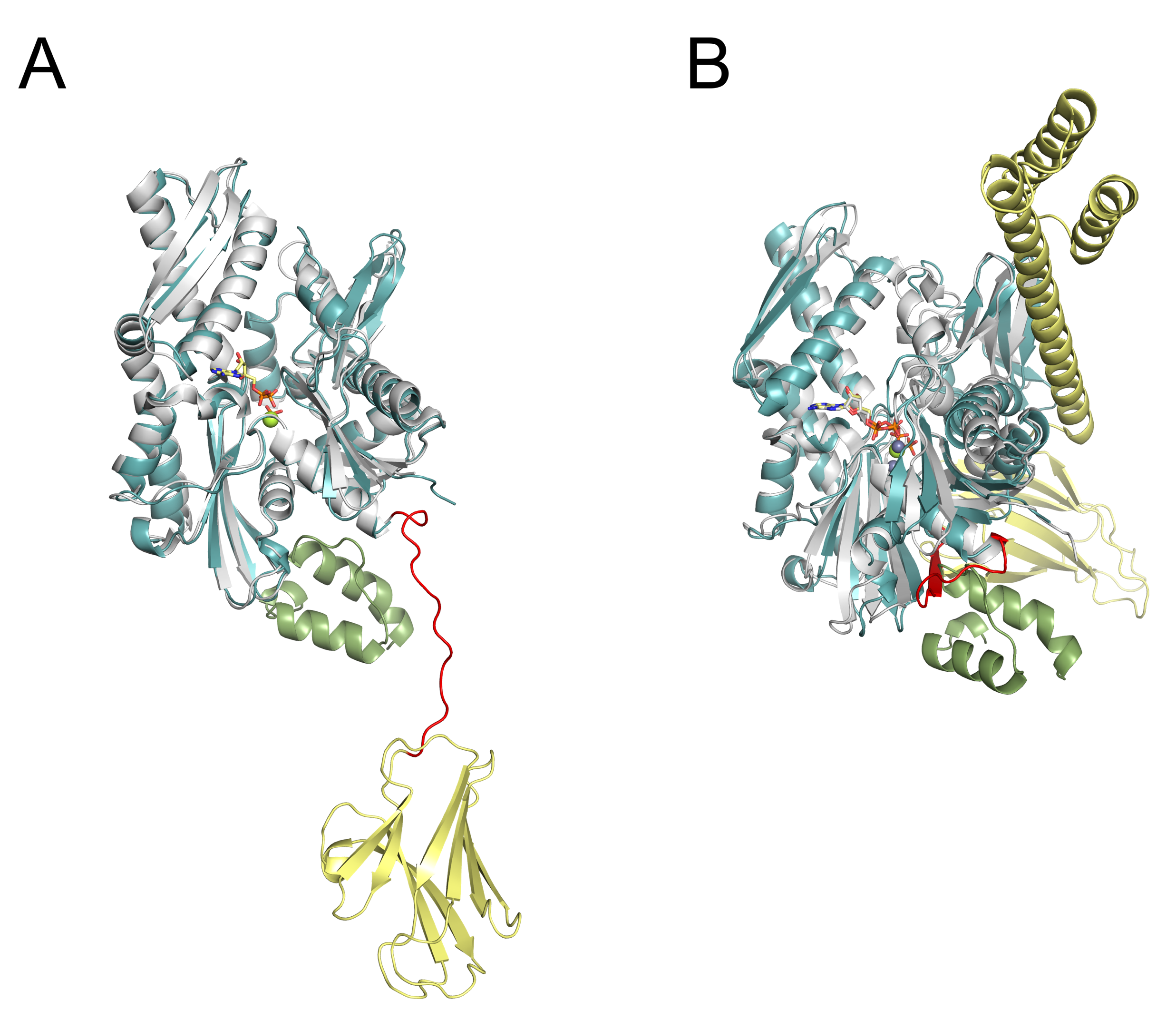

**Figure S3. CDNF binds BiP in cells as shown by BiFC, while selected CDNF mutants show reduced or altered interaction.** Positive control (Jun-NVenus + Fos-CVenus) and negative control (Jun-NVenus + Max-CVenus) demonstrate assay specificity. Wild-type CDNF and BiP show a strong BiFC signal, indicating interaction. CDNF mutants (V118Y, A134Q, and E137A) display altered levels of BiFC signal with BiP. A yellow fluorescent signal indicates interaction between the tested proteins. The endoplasmic reticulum (ER) was visualized using ER-ID-red dye (red), and nuclei were stained with Hoechst 33342 (blue). Scale bars: 10 µm.

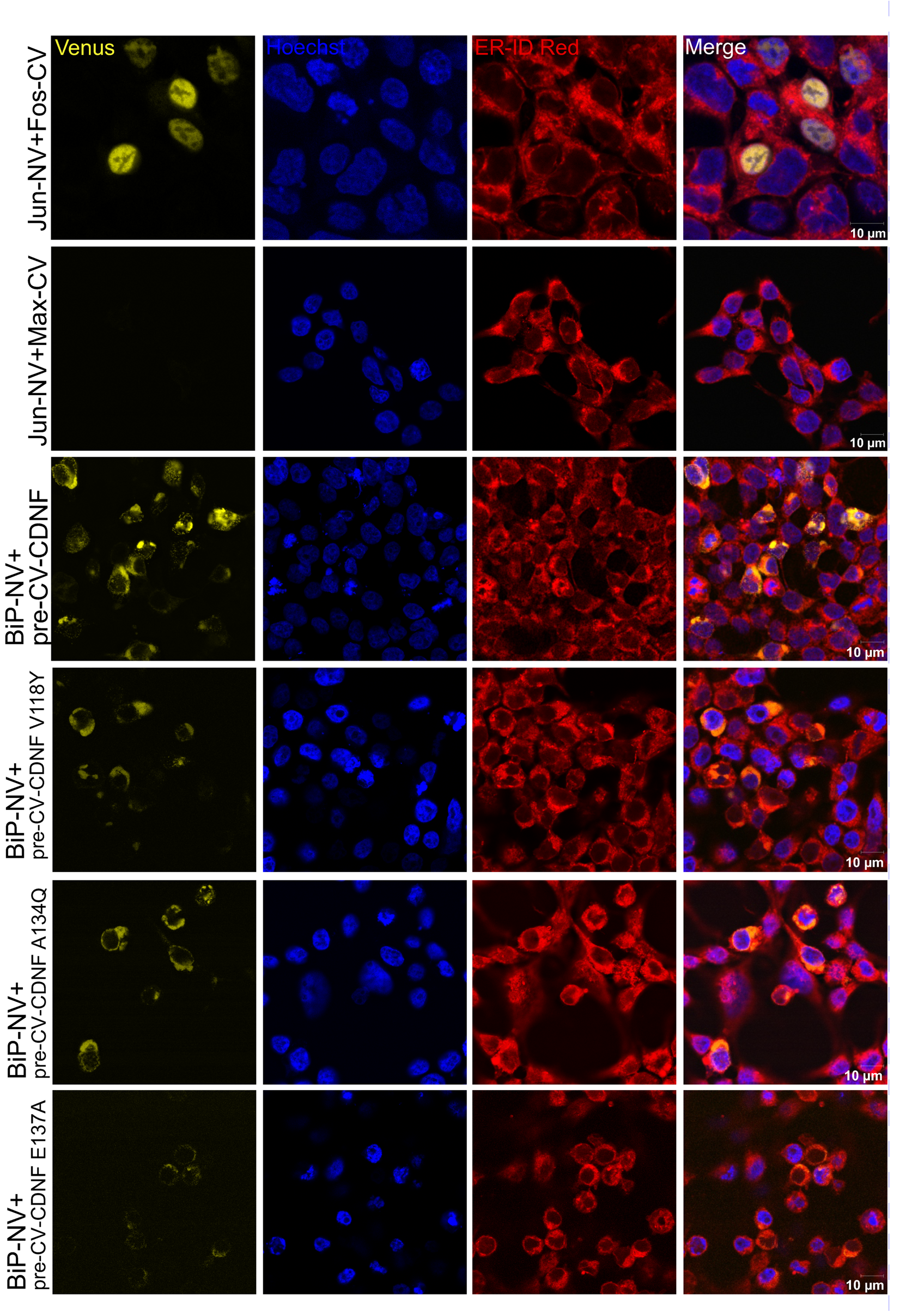

**10 µm**

**Figure S4.** Analysis of intermolecular contacts between BiP NBD and C-CDNF in our MD simulation trajectories of the high-resolution apostructure of the NBD-CDNF complex. The graph shows all pairs of residues which (on average) create more than 40 contacts/frame throughout the trajectory (cutoff value of 5 Å, see methods).

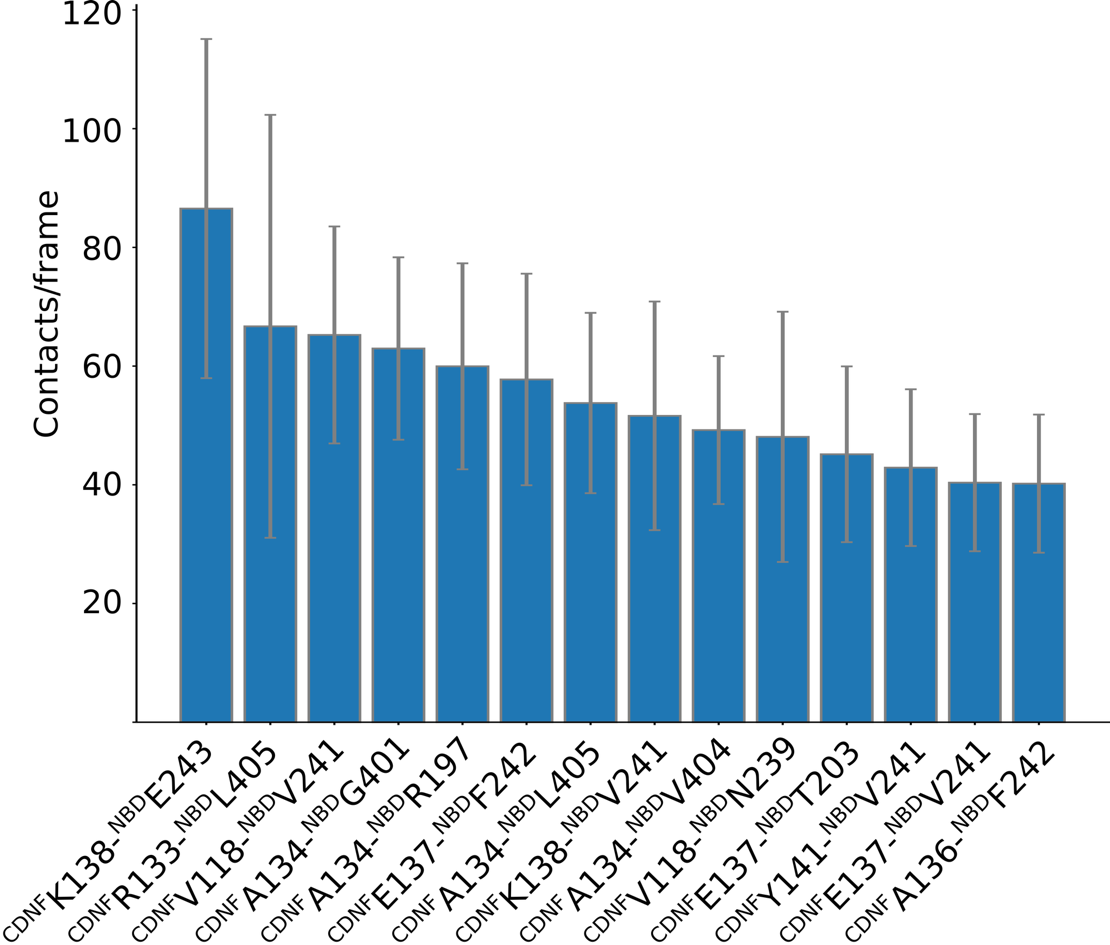

**Figure S5. Molecular dynamics simulations trajectories.** (**A**) Root-mean-square deviation (RMSD) of C_α_ carbon atoms of BiP NBD in a free state (right panel) and in complex with C-CDNF (left panel). (**B**) Root-mean-square fluctuation (RMSF) of C_α_ carbons of BIP NBD in complex with C-CDNF (in blue), compared to the free NBD (in red). The error bars in (**B**) show the standard deviation of RMSD values along the trajectory.

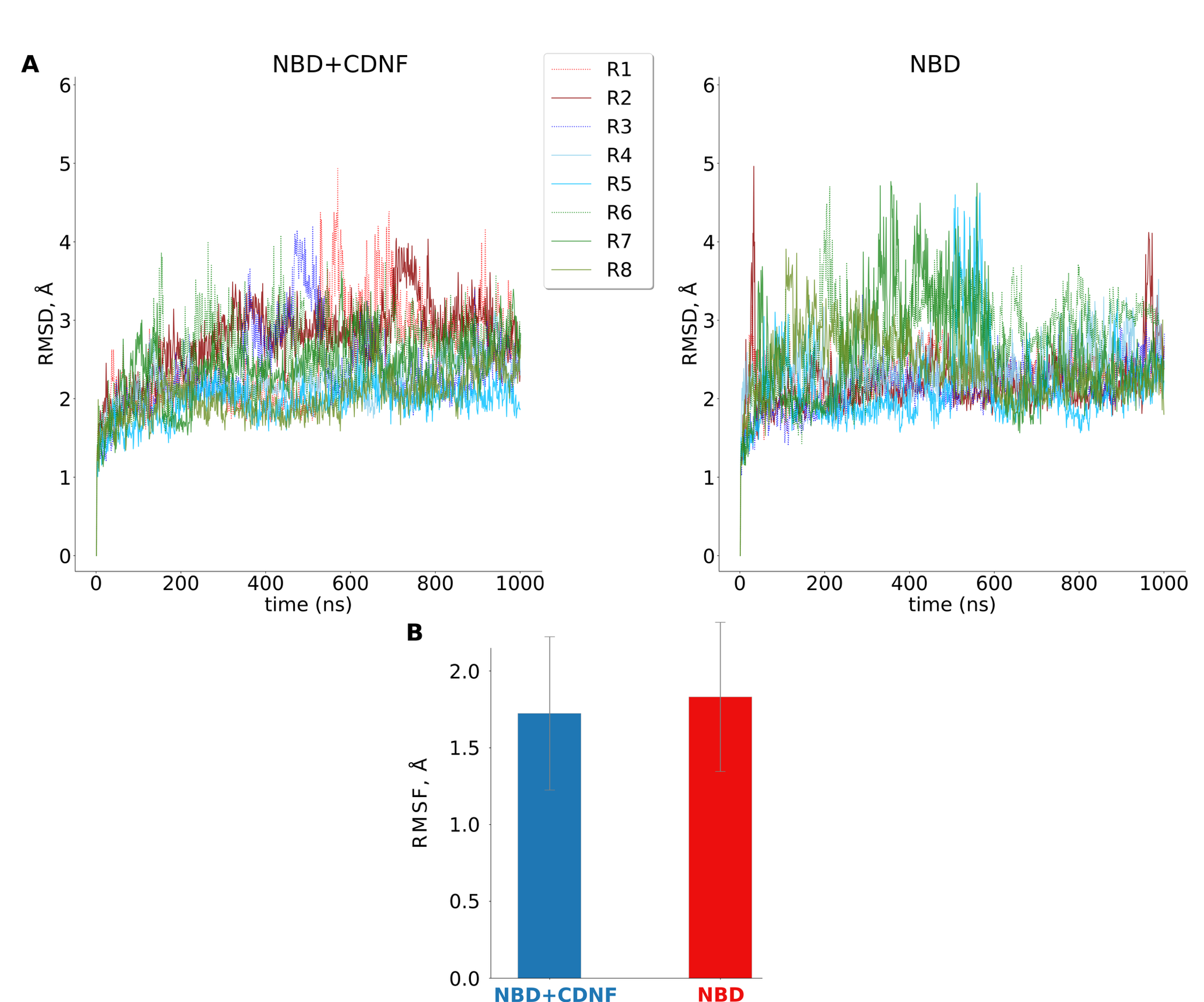

**Figure S6**. Notation of lobes in the C-CDNF-bound structure of the BiP nucleotide-binding domain, with the location of the residue E293 and K81 forming a salt bridge over the nucleotide binding site indicated.

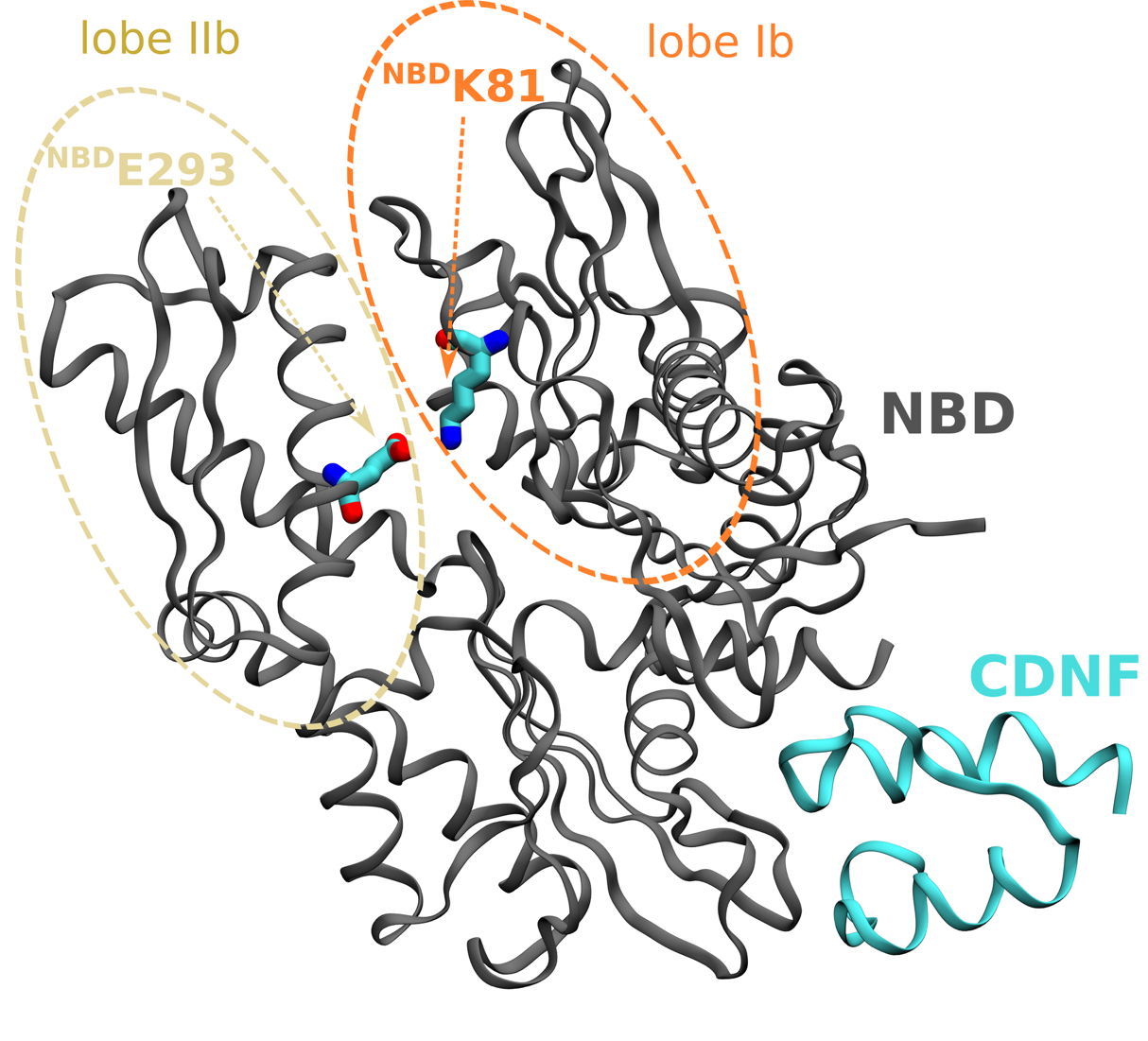

**Figure. S7.** Time series of the distances between ADP and ^NBD^Ser365 in the system with free BiP NBD (right curves, red), and BiP NBD bound to CDNF (left curves, blue). The distance is calculated between the ADP purine ring (atoms N1-C2-N3-C4-C5-C6) and the Ca carbon of ^NBD^Ser365 (see methods, and Fig. 5D).

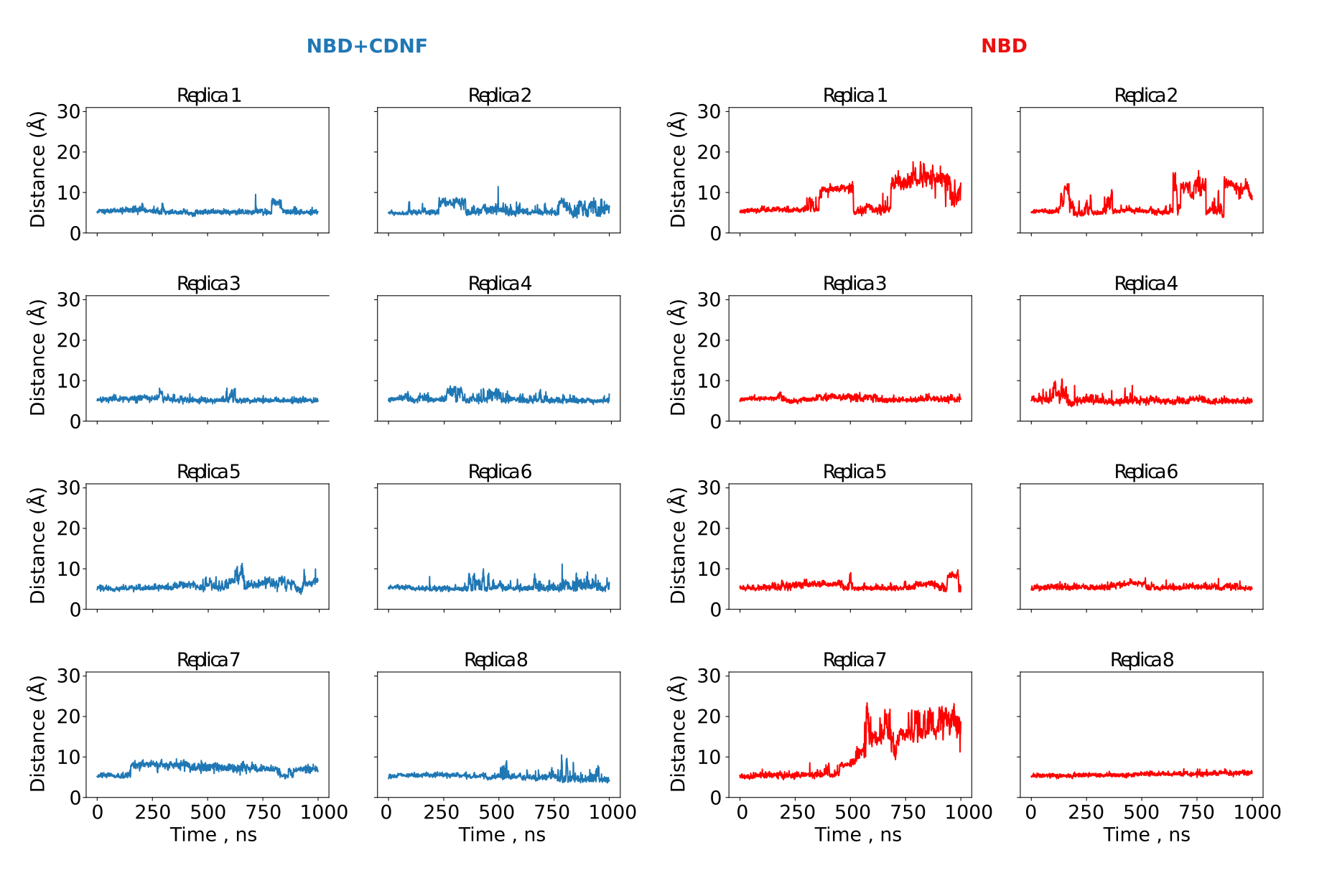

**Figure S8. Comparison of yeast Sse1 and C-CDNF binding to Hsp70/BiP NBD**. The yeast NEF homolog of Grp170 (Sse1) structure (PDB: 3D2F) in the complex with Hsp70 NBD (Sse1, green, NBD cyan) compared to BiP NBD conformation in presence of CDNF. Note the dramatic movement of lobe IIb (arrow, ca. 27° rotation) due to binding of Sse1, its mammalian homolog Grp170 is presumed to has the same effect as Sse1, given the they are functionally homologous. The CNDF bound BiP NBD is shown in red. C-CDNF labeled (blue).

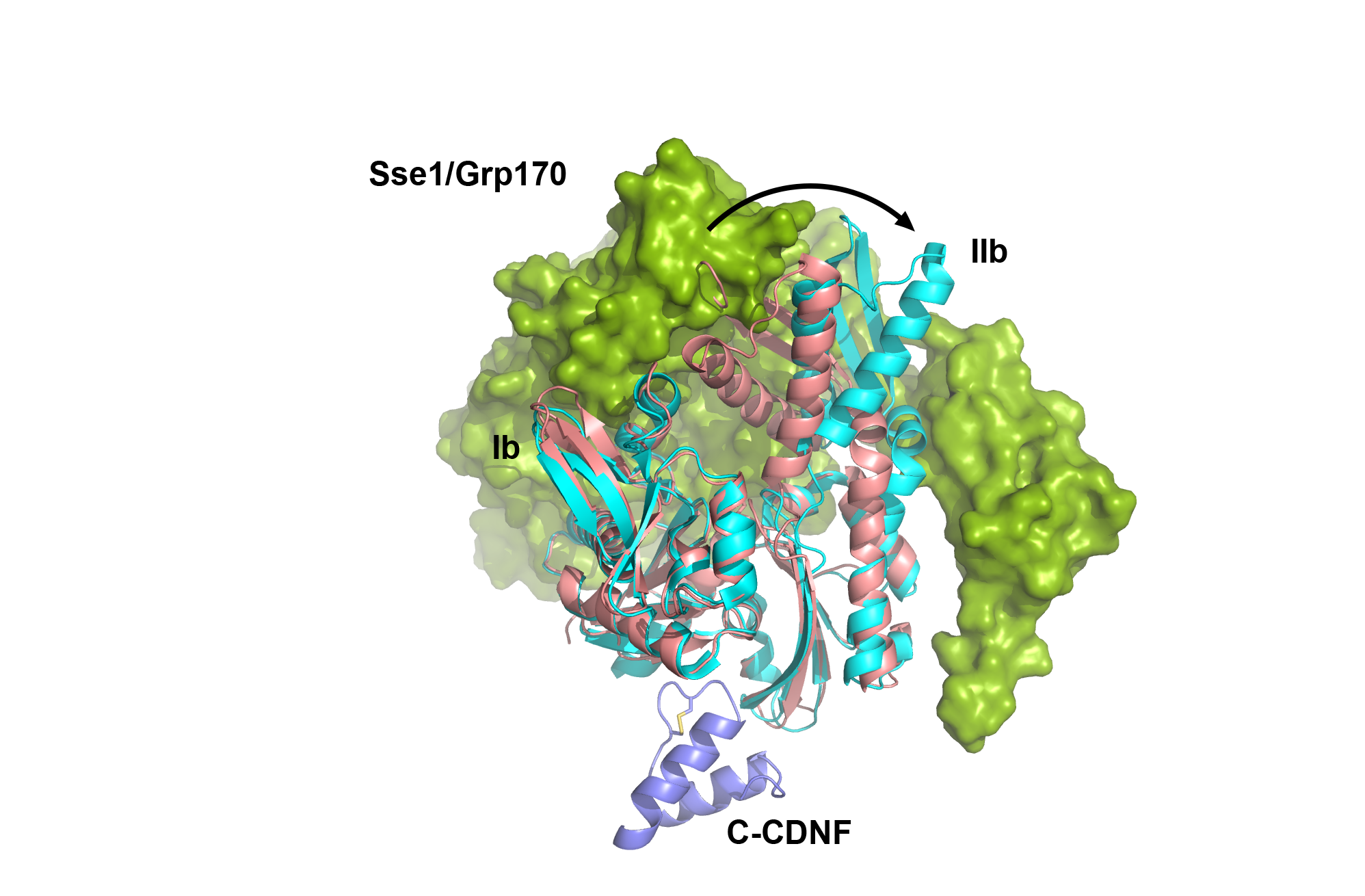

**Supplementary Table 1.** Crystallographic data collection and refinement

|  | BiP NBD & C-CDNF | BiP NBD & C-CDNF & ADP |
| --- | --- | --- |
| **Data collection** |  |  |
| Beamline | DLS I24 | ESRF ID30A-3 |
| Wavelength (Å) | 0.99987 | 0.96770 |
| Space group | *P*2_1_2_1_2_1_ | *P*2_1_2_1_2_1_ |
| Cell dimentions |  |  |
| *a*, *b*, *c* (Å) | 50.52, 80.61, 102.35 | 50.63, 80.94, 102.99 |
| α, β, γ (°) | 90, 90, 90 | 90, 90, 90 |
| Resolution (Å) | 40.31-1.65 (1.68-1.65) | 51.49-1.50 (1.53-1.50) |
| No. of reflections | 215747 (8357) | 859226 (23990) |
| No. of unique reflections | 50649 (2315) | 68155 (3180) |
| *R*_merge_ | 0.108 (1.129) | 0.129 (1.577) |
| I/σ(I) | 8.11 (1.26) | 13.2 (1.4) |
| CC_1/2_ | 0.995 (0.431) | 0.999 (0.471) |
| Completeness (%) | 99.2 (92.4) | 99.5 (94.7) |
| Multiplicity | 4.3 (3.6) | 12.6 (7.5) |
| **Refinement** |  |  |
| *R*_work_/*R*_free_ | 0.181/0.203 | 0.176/0.198 |
| No. of atoms |  |  |
| Protein | 3275 | 3302 |
| Water | 361 | 542 |
| Others | 13 | 33 |
| *B*-factor (Å^2^) |  |  |
| Protein | 25.5 | 20.4 |
| Water | 32.9 | 31.4 |
| Others | 26.1 | 16.3 |
| RMS bond length (Å) | 0.009 | 0.004 |
| RMS bond angle (°) | 1.464 | 0.805 |
| Ramachandran favored (%) | 99.5 | 100.0 |
| Ramachandran outliers (%) | 0 | 0 |

Values in parenthesis refer to the highest resolution shell.

**Supplementary Table 2.** Buried surface area for interface residues at the CNDF-BiP binding interface.

| **CDNF-BiP complex (Interface area: 558.5 Å^2^)** | | |  |
| --- | --- | --- | --- |
| **CDNF** | **Solvent accessible area (Å**^2^) | **Buried surface area (Å**^2^) | **% buried surface area** |
| VAL 118 | 67.56 | 64.09 | 94.87 |
| ALA 136 | 100.91 | 91.37 | 90.54 |
| ALA 134 | 108.48 | 97.10 | 89.52 |
| GLU 137 | 117.43 | 92.39 | 78.68 |
| LYS 138 | 74.56 | 43.54 | 58.40 |
| **BiP** | **Solvent accessible area (Å**^2^) | **Buried surface area (Å**^2^) | **% buried surface area** |
| VAL 241 | 103.59 | 99.74 | 96.29 |
| THR 203 | 20.74 | 20.58 | 99.20 |
| PHE 242 | 36.47 | 35.22 | 96.55 |
| GLN 401 | 60.76 | 44.16 | 72.68 |
| ASP 238 | 86.17 | 61.67 | 71.57 |
| ASN 200 | 33.55 | 18.56 | 55.31 |
| ASN 239 | 114.77 | 64.21 | 55.95 |
| **CDNF-BiP-ADP complex (Interface area: 564.8 Å^2^)** | | |  |
| **CDNF** | **Solvent accessible area (Å**^2^) | **Buried surface area (Å**^2^) | **% buried surface area** |
| VAL 118 | 65.45 | 62.46 | 95.42 |
| ALA 134 | 106.09 | 91.43 | 86.19 |
| ALA 136 | 102.25 | 91.14 | 89.13 |
| GLU 137 | 115.45 | 94.42 | 81.79 |
| LYS 138 | 75.32 | 45.01 | 59.76 |
| **BiP** | **Solvent accessible area (Å**^2^) | **Buried surface area (Å**^2^) | **% buried surface area** |
| VAL 241 | 103.92 | 100.20 | 96.42 |
| THR 203 | 20.68 | 20.52 | 99.20 |
| PHE 242 | 35.26 | 33.85 | 95.99 |
| ASP 238 | 85.25 | 61.94 | 72.66 |
| ILE 207 | 11.83 | 7.70 | 65.12 |
| GLN 401 | 56.53 | 39.05 | 69.07 |
| ASN 200 | 35.92 | 19.46 | 54.16 |
| ASN 239 | 122.64 | 70.97 | 57.86 |

**Supplementary Table 3**. Hydrogen bonded interactions at the CNDF-BiP binding interface.

|  | CDNF-BiP complex |  |
| --- | --- | --- |
| **CDNF** | **Distance (Å)** | **BiP** |
| ARG 117 [NE] | 3.10 | ASP 238 [OD2] |
| ARG 117 [NH2] | 3.04 | ASP 238 [OD1] |
| VAL 118 [N] | 3.38 | ASP 238 [OD2] |
| LYS 122 [NZ] | 2.77 | ASN 239 [O] |
| ARG 133 [NH1] | 2.67 | VAL 404 [O] |
| LYS 138 [N] | 3.01 | PHE 242 [O] |
| ARG 133 [O] | 3.01 | ARG 197 [NH2] |
| ALA 134 [O] | 3.13 | ARG 197 [NE] |
| ALA 134 [O] | 3.37 | GLN 401 [NE2] |
| ALA 136 [O] | 3.04 | PHE 242 [N] |
| GLU 137 [OE2] | 2.85 | THR 203 [OG1] |
| GLU 137 [OE2] | 2.80 | ASN 200 [ND2] |

|  | CDNF-BiP-ADP complex |  |
| --- | --- | --- |
| **CDNF** | **Distance (Å)** | **BiP** |
| ARG 117 [NE] | 2.96 | ASP 238 [OD2] |
| ARG 117 [NH2] | 3.03 | ASP 238 [OD1] |
| VAL 118 [N] | 3.34 | ASP 238 [OD2] |
| LYS 122 [NZ] | 2.73 | ASN 239 [O] |
| ARG 133 [NH1] | 2.56 | VAL 404 [O] |
| LYS 138 [N] | 2.94 | PHE 242 [O] |
| ARG 133 [O] | 2.86 | ARG 197 [NH2] |
| ALA 134 [O] | 3.16 | ARG 197 [NE] |
| ALA 136 [O] | 3.04 | PHE 242 [N] |
| GLU 137 [OE2] | 2.75 | THR 203 [OG1] |
| GLU 137 [OE2] | 2.83 | ASN 200 [ND2] |
